# 3’UTR-directed, kinase proximal mRNA decay inhibits C/EBPβ phosphorylation/activation to suppress senescence in tumor cells

**DOI:** 10.1101/2022.03.29.486281

**Authors:** Jacqueline Salotti, Nida Asif, Srikanta Basu, Aniruddha Das, Mei Yang, Baktiar Karim, Karen Saylor, Nancy Martin, David A. Scheiblin, Sweta Misra, Brian T. Luke, Thorkell Andresson, Stephen Lockett, Lino Tessarollo, Peter F. Johnson

## Abstract

C/EBPβ is a potent regulator of oncogene-induced senescence (OIS) and the SASP. C/EBPβ is post-translationally activated in OIS cells by the effector kinases ERK1/2 and CK2. However, in tumor cells C/EBPβ activation is suppressed by its 3’UTR. 3′UTR regulation of protein activity (UPA) requires a G/U-rich element (GRE) and its cognate binding protein, HuR. These components segregate *CEBPB* transcripts away from a perinuclear compartment harboring ERK1/2 and CK2, restricting C/EBPβ from its activating kinases. We report here that the mRNA decay proteins UPF1 and Staufen1/2 are essential UPA factors enriched within the perinuclear cytoplasm. STAU1/2 and UPF1 overlap with CK2 on perinuclear signaling endosomes where they promote localized *CEBPB* mRNA decay. UPF1 or STAU1/2 depletion in tumor cells increased *CEBPB* transcripts adjacent to CK2 foci, coinciding with C/EBPβ activation and senescence. The GRE and an adjacent STAU binding site independently suppress C/EBPβ-mediated senescence, while a distinct 3’UTR region inhibits its SASP-inducing activity. *Kras^G12D^*-driven lung tumors in mice carrying a *Cebpb* GRE deletion rarely progressed to malignant adenocarcinomas, demonstrating the importance of UPA to enable tumor progression *in vivo*. Thus, kinase-proximal mRNA decay is a novel mechanism that inhibits C/EBPβ activation in tumor cells to facilitate senescence bypass.

## Introduction

Senescence is a persistent state of cell cycle arrest elicited by stress stimuli such as DNA damage, oxidative stress and oncogenes.^1^ Oncogene-induced senescence (OIS) triggered by activated *RAS*, *BRAF*, *MYC* or other oncogenes establishes an intrinsic barrier to neoplastic transformation and cancer.^2^ The OIS checkpoint imposes selective pressure for mutations or epigenetic events that disable key pro-senescence pathways such as ARF-p53 and p16-RB, allowing premalignant cells to bypass senescence and progress to a fully transformed state. Senescent cells also express a characteristic set of secreted proteins termed the senescence-associated secretory phenotype (SASP)^3^ primarily composed of pro-inflammatory cytokines, their receptors and matrix proteases. SASP factors reinforce and propagate senescence through autocrine and paracrine mechanisms.^4–6^ SASP cytokines released by senescent cells also mediate recognition and clearance by the immune system (senescence surveillance), which is critical for efficient tumor suppression *in vivo*.^7^

NF-κB and C/EBPβ are key transcriptional regulators of senescence and the SASP.^4,8–11^ Tumor cells typically display relatively low expression of SASP genes. This “cold” inflammatory profile may contribute to the cells’ ability to evade immune-surveillance. Paradoxically, many cancer cells express C/EBPβ and activated NF-κB, which would be expected to induce senescence and the SASP. However, the pro-senescence function of C/EBPβ is inhibited in tumor cells by a 3’UTR-dependent mechanism that suppresses RAS-induced post-translational activating modifications.^12,13^ The 3’UTR also potently inhibits the transcriptional activity of C/EBPβ toward SASP genes. This mechanism (3’UTR regulation of protein activity, or UPA) involves subcellular partitioning of *Cebpb* transcripts, excluding them from a perinuclear cytoplasmic region containing the C/EBPβ kinases, ERK1/2 and CK2. C/EBPβ translated in this location is unable to access its activating kinases, suppressing phosphorylation on residues Thr188 (ERK1/2) and Ser222 (CK2) and possibly other sites.^12,13^ This hypo-phosphorylated state restrains C/EBPβ DNA binding, homodimerization and transcriptional potential, which underlie its cytostatic and SASP-inducing activities in senescent cells.^12,14–16^

UPA is disabled in primary cells undergoing oncogenic RAS-induced senescence (RIS), allowing C/EBPβ activation by senescence stimuli. UPA is inactivated by a RAS-CaMKKβ-AMPK pathway that induces nuclear translocation of HuR/ELAVL1^17^, an RNA-binding protein that recognizes a G/U-rich element (GRE) region in the *Cebpb* 3’UTR.^13,18^ HuR and the GRE motif are required for C/EBPβ UPA in tumor cells, as HuR ablation or deleting the GRE abrogates 3’UTR inhibition and disrupts peripheral *CEBPB* mRNA localization.^13^ Cytoplasmic HuR levels are elevated in tumor cells and promote increased tumor malignancy^19^ by facilitating unrestrained proliferation and senescence bypass through various target mRNAs, including repressing C/EBPβ through the UPA mechanism.

The role of HuR in C/EBPβ UPA and the mechanism by which the 3’UTR controls *Cebpb* mRNA localization are important but unresolved questions. Since HuR alone is unlikely to control C/EBPβ UPA, we sought to characterize other components of the UPA machinery and gain further insights into the underlying mechanism.

## Results

### Identification of *Cebpb* GRE-associated proteins

We hypothesized that a higher-order protein complex assembles on the GRE:HuR scaffold to instate UPA and inhibit C/EBPβ activity (Fig. 1A). To identify additional *Cebpb* GRE-interacting factors, we used an *in vitro* RNA affinity purification/mass spectrometry (MS) procedure previously employed to characterize proteins that bind to A/U-rich elements (AREs)^20^ (Fig. S1A). Proteins associating with the 102 nt *Cebpb* GRE sequence were purified from NIH 3T3 cell cytoplasmic lysates, as these immortalized fibroblasts are UPA proficient and therefore express the relevant factors. Specific enrichment was determined by comparing GRE-bound proteins to those interacting with a control GFP RNA of similar length. The GRE-bound fraction revealed ∼50 visible bands on a Coomassie stained protein gel (Fig. 1B). Proteins from three independent pull-down experiments were analyzed by mass spectrometry, with GRE binders distinguished from non-specific background by calculating GRE:GFP peptide ratios. Using a threshold of log_2_ GRE:GFP ratio ≥1; -log_10_ FDR ≥1, 324 GRE-interacting proteins were identified (Fig. 1C; Table S1). The GRE-bound list was analyzed by gene ontology (DAVID, GOTERM_BP)^21^ to identify enriched biological processes (Fig. 1D; Table S1). The categories were skewed toward RNA-associated functions such as translation, splicing and RNA decay.

**Figure 1.**
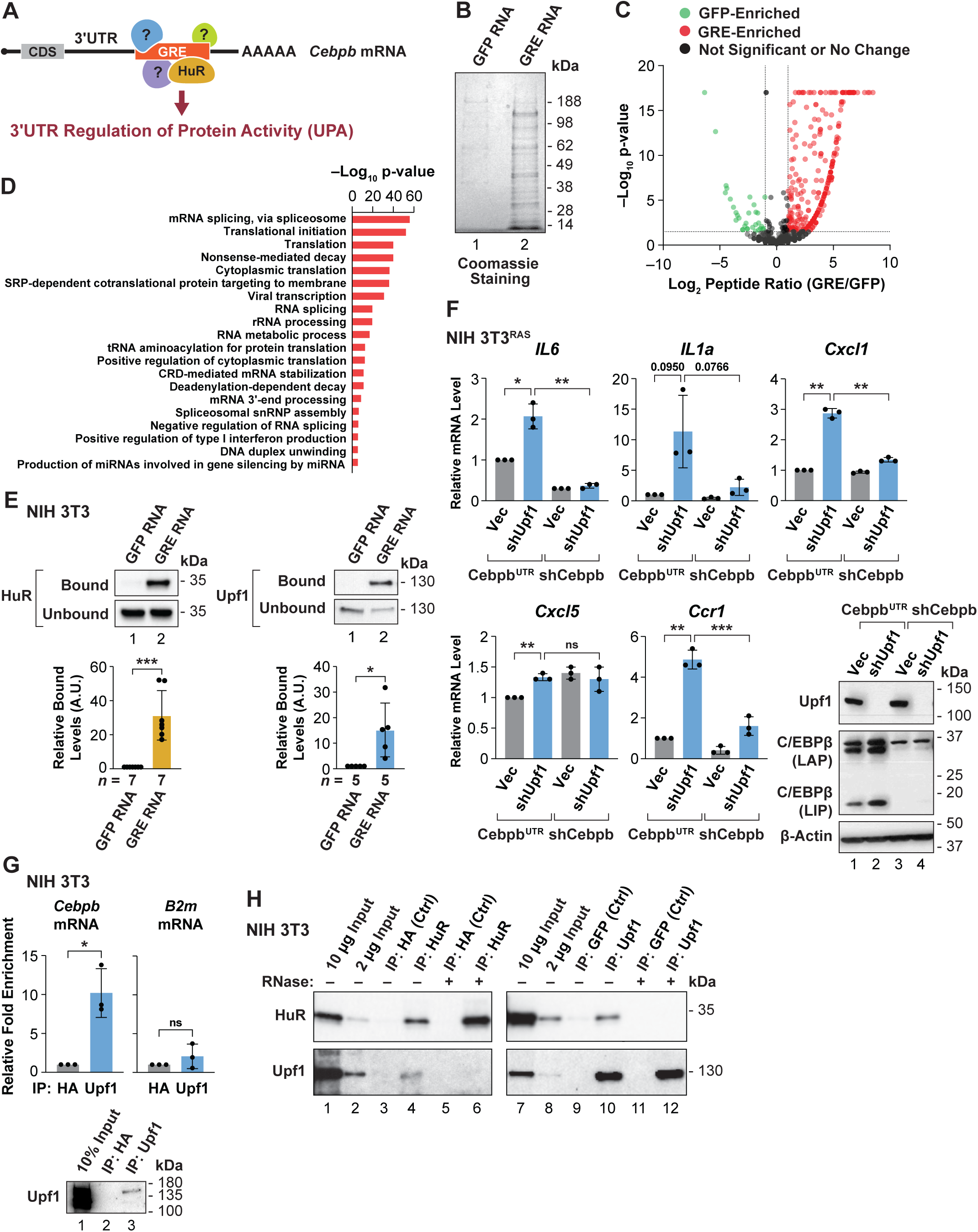
The *Cebpb* 3’UTR GRE motif associates with mRNA decay factor Upf1. (**A)** Model for binding of HuR and other proteins to the *Cebpb* GRE to establish 3’UTR inhibition (UPA). **(B)** Coomassie stained gel of proteins binding to the GRE RNA or a control GFP RNA. **(C)** Volcano plot of proteins displaying positive or negative enrichment for GRE binding, as determined by mass spectrometry. GRE/GFP peptide count ratios are the sum of three independent affinity purification experiments. Thresholds of log_2_ GRE:GFP ratio ≥ 1; -log_10_ FDR ≥ 1 were used to identify 324 GRE-interacting proteins. Forty-four proteins had p-values < 10^−17^ and were plotted as equal to 10^−17^ for graphing purposes only. **(D)** Gene ontology analysis of proteins preferentially binding to the GRE. GRE-enriched proteins were analyzed for associations with pathways/biological processes using DAVID ^21^. GO terms for the highest-ranking biological processes are shown, ranked by P-value. **(E)** Immunoblots confirming binding of HuR and Upf1 to the GRE RNA. Quantitation of binding enrichment (average of 5 experiments) is shown below. **(F)** Upf1 was ablated in 3T3^RAS^ cells overexpressing *Cebpb^UTR^*(βUTR) or depleted for C/EBPβ (shβ) and expression of selected proinflammatory SASP mRNAs was analyzed by RT-qPCR. **(G)** Association of Upf1 with *Cebpb* or *B2m* (control) transcripts in NIH 3T3 cells. nRIP assays from 3 independent assays are shown. Bottom: immunoblots showing IP of Upf1. **(H)** HuR and Upf1 associate in an RNA-dependent manner. IPs were performed using NIH 3T3 cell lysates in the absence or presence of 200 µg/mL RNase A. Statistical significance was determined using Student’s t test. *p < 0.05,**p < 0.01,***p < 0.001.

As a secondary screen for functional UPA proteins, we used RNAi to deplete a set of 21 prominent GRE interactors (ranked by peptide ratios), as well as other candidates identified in a pilot GRE pull-down experiment (Table S1). We then examined expression of several C/EBPβ-regulated SASP genes, as depletion of bona fide UPA factors should activate C/EBPβ and induce the SASP. These tests were performed in HRAS^G12V^-tranformed NIH 3T3 cells transduced with a *Cebpb*^UTR^ construct (3T3^RAS^-β^UTR^), since NIH 3T3^RAS^ cells express low levels of endogenous C/EBPβ.^22^ The same knockdowns were performed in 3T3^RAS^ cells in which endogenous C/EBPβ was further depleted to demonstrate C/EBPβ dependence. As shown in Fig. S1B, *Il6, Il1α, Cxcl1* and *Ccr1* mRNAs were increased upon knockdown of Strbp and Upf1 (RENT1) in 3T3^RAS^-β^UTR^ cells but not in 3T3^RAS^-sh*Cebpb* cells. Silencing of other candidates such as Elavl1 (HuR) and Fubp3 also induced *Ccr1* expression but did not appreciably activate the other SASPs. These data implicate Upf1 and Strbp as potential components of the UPA system.

Upf1 is an RNA helicase/ATPase and RNP remodeling protein that is a key regulator of nonsense-mediated mRNA decay (NMD) and other RNA degradation pathways.^23^ We further investigated the role of Upf1 in C/EBPβ UPA since its depletion caused the largest induction of SASP genes (Fig. S1B) and UPF1 association with *CEBPB* mRNA is seen in genome-wide CLIP-seq studies^24,25^. In addition, a database of RNA-independent and -dependent Upf1-interacting proteins^26^ overlapped considerably with our list of GRE binders (Fig. S1C; Table S2). UPF1 ablation also displays synthetic lethality with oncogenic KRAS in tumor cells^27^, consistent with our prediction that disruption of C/EBPβ UPA impairs proliferation/survival of RAS-transformed cells. Immunoblotting verified selective enrichment of Upf1 and HuR in the GRE-bound fraction compared to the control RNA (Fig. 1E).

We confirmed that Upf1 depletion activates SASP genes in a C/EBPβ-dependent manner in 3T3^RAS^ cells (Fig. 1F). Native RNA immunoprecipitation (nRIP) demonstrated that Upf1 associates with *Cebpb* transcripts but not a control gene (β_2_ microglobulin) (Fig. 1G). To test whether Upf1 and HuR interact, we performed co-immunoprecipitation assays using NIH 3T3 cell lysates (Fig. 1H). HuR and Upf1 displayed reciprocal co-immunoprecipitation and this interaction was disrupted by addition of RNase, indicating that the two proteins associate in an RNA-dependent manner. These findings are consistent with proteomic data identifying HuR as an RNA-dependent UPF1 interactor.^26^

### Perinuclear Upf1-dependent mRNA decay controls subcellular partitioning of *CEBPB* transcripts

To investigate if Upf1 contributes to UPA-mediated inhibition of C/EBPβ activity, we first examined the subcellular distribution of *Cebpb* transcripts and Upf1 in NIH 3T3 cells. RNA FISH showed that *Cebpb* mRNAs were excluded from the perinuclear region (Fig. S2A), as observed previously.^17^ Moreover, Upf1 was restricted to the perinuclear area that was largely devoid of *Cebpb* FISH signals. When Upf1 was depleted, *Cebpb* transcripts became more abundant in this region. This effect was particularly evident in individual cells displaying efficient knockdown of Upf1.

NIH 3T3 cells are not transformed and hence do not display constitutive perinuclear localization of CK2 and p-ERK^12^, and the elongated spindle morphology of RAS-transformed NIH 3T3 cells is not conducive to subcellular localization analysis. Therefore, to investigate the spatial relationship between *CEBPB* mRNA, UPF1 and kinases we analyzed human A549 lung adenocarcinoma cells (*KRAS^G12S^*). Immunofluorescence imaging showed that UPF1 was perinuclear in these cells, as was CK2α, and both proteins were largely non-overlapping with *CEBPB* transcripts (Fig. 2A). UPF1 depletion increased the abundance of nuclear proximal *CEBPB* transcripts, which was confirmed by measuring the relative nuclear proximity index (RNPI)^28^ for *CEBPB* RNA FISH signals in control and UPF1-depleted cells (Fig. 2A). To analyze overlap with CK2, we used CK2α signal thresholding to define a perinuclear kinase region and determined the percentage of *CEBPB* FISH signals present within the CK2 boundary (Fig. 2B). mRNA-CK2 overlap averaged 25.6% in control cells but increased to 62.6% when UPF1 was depleted (p<0.0001).

**Figure 2.**
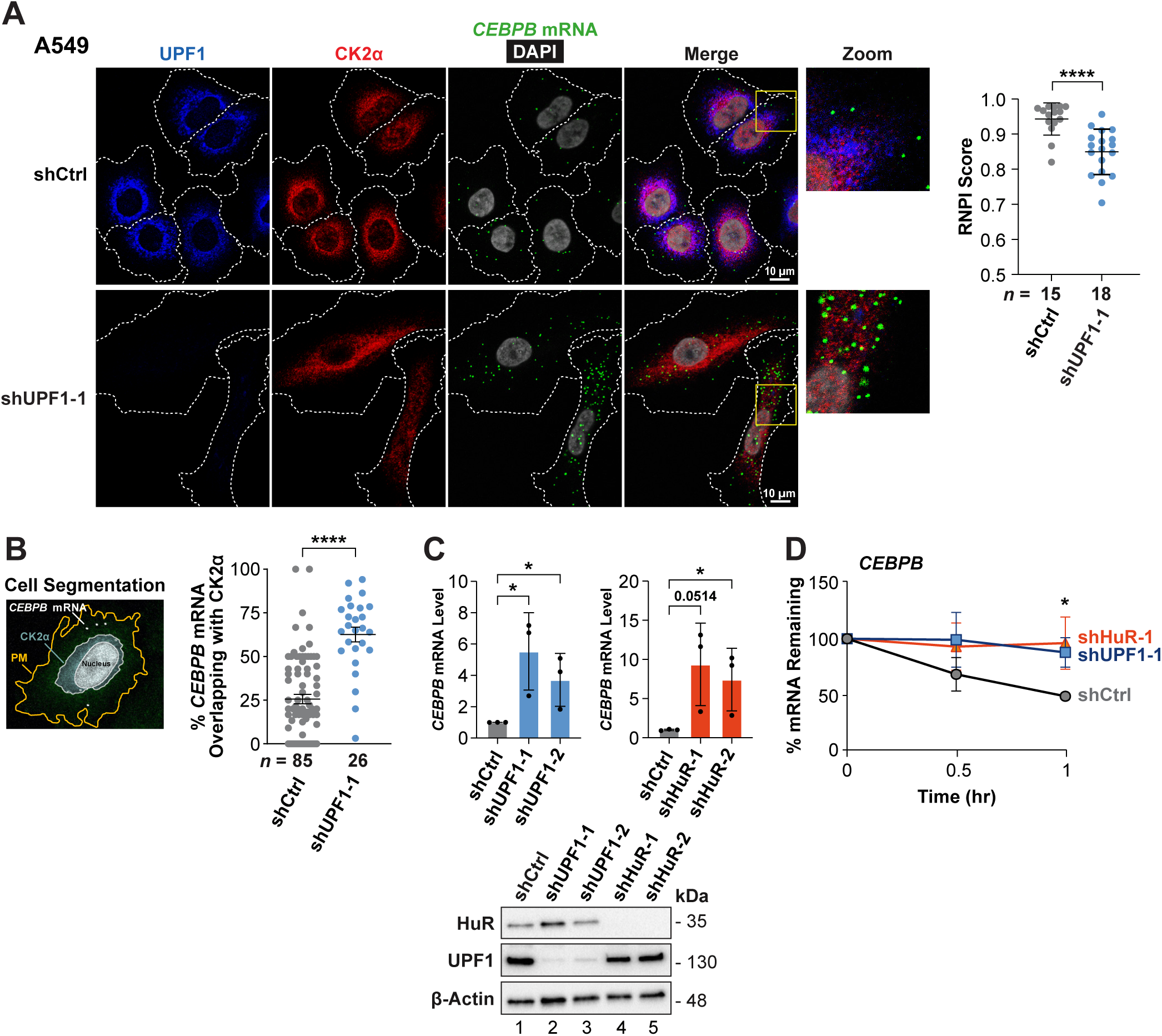
UPF1 controls perinuclear decay of *CEBPB* transcripts. **(A)** UPF1 depletion in A549 lung adenocarcinoma cells increases the abundance of *CEBPB* transcripts in the perinuclear region containing CK2. *CEBPB* mRNA was detected using single molecule RNA FISH in combination with UPF1 and CK2α IF staining. Nuclear proximity of *CEBPB* transcripts (RNPI score/cell) is shown on the right. **(B)** Overlap of perinuclear CK2α and *CEBPB* mRNA increases after UPF1 silencing in A549 cells. A perinuclear CK2α domain was defined by fluorescence thresholding and the number of cytoplasmic *CEBPB* mRNA signals within this domain was determined for each cell. *n* = number of cells analyzed. **(C)** UPF1 or HuR depletion increases total *CEBPB* mRNA levels in A549 cells. mRNA levels were determined by RT-qPCR analysis of RNA samples, normalized to *PPIA* mRNA. **(D)** *CEBPB* mRNA is stabilized by depletion of UPF1 or HuR. Decay rates were measured by EU pulse-chase labeling, followed by RT-qPCR analysis of EU-containing *CEBPB* transcripts, normalized to *PPIA* mRNA. *n* = number of cells analyzed. Statistical significance was determined using Student’s t test. *p < 0.05,****p < 0.0001.

As UPF1 plays a key role in mRNA decay, its restriction to perinuclear areas that are devoid of *CEBPB* transcripts suggested that localized degradation may prevent mRNA accumulation in these regions. Supporting this idea, depletion of UPF1 or HuR in A549 cells increased the overall abundance of *CEBPB* mRNAs, as determined by RT-qPCR assays (Fig. 2C). Similar results were obtained with RAS-transformed NIH 3T3 cells (Fig. S2B). Accordingly, UPF1 depletion was associated with increased *CEBPB* mRNA half-life (Fig. 2D). *CEBPB* mRNA stabilization was also observed upon HuR silencing, which causes increased perinuclear localization of *CEBPB* transcripts.^13^ These observations indicate that UPF1 and HuR promote perinuclear mRNA decay (PMD).

### UPF1 suppresses C/EBPβ phosphorylation and senescence in tumor cells

The segregation of *CEBPB* transcripts from perinuclear CK2 in tumor cells inhibits C/EBPβ phosphorylation on a CK2 site (Ser222 in mouse, Ser271 in human), which stimulates C/EBPβ DNA binding and its pro-senescence activity.^12^ Super-resolution imaging of A549 cells depleted for UPF1 revealed many *CEBPB* transcripts located adjacent to CK2α foci (Fig. 3A), which correspond to perinuclear signaling endosomes.^12,28^ CK2α-*CEBPB* mRNA distance relationships were quantified by nearest neighbor image analysis (16 cells/group), revealing significantly increased proximity upon UPF1 depletion (Fig. 3A, right panel). Proximity labeling assays (PLA) using C/EBPβ and CK2α antibodies likewise showed increased cytoplasmic contiguity of the two proteins in UPF1 knockdown cells compared to controls (Fig. 3B). Thus, when UPF1 is depleted C/EBPβ proteins occur more frequently at perinuclear CK2 signaling endosomes, prior to nuclear translocation. This suggests that when UPA-dependent perinuclear mRNA decay is interrupted, C/EBPβ phosphorylation by CK2 is facilitated as *CEBPB* transcripts access signaling endosomes. Since we were unable to generate an antibody that efficiently detects human C/EBPβ p-Ser271, we used a p-Ser222 antibody to monitor phosphorylation of mouse C/EBPβ in the chromatin fraction of 3T3^RAS^-β^UTR^ cells. UPF1 depletion augmented p-Ser222 levels (normalized to total C/EBPβ) by 4.6-fold (Fig. 3C) and increased C/EBPβ DNA binding and homodimerization, with the proportion of EMSA homodimer species elevated 8-fold compared to the control (Fig. S3A). Enhanced C/EBPβ DNA binding was also observed upon UPF1 knockdown in PANC-1 PDAC cells and HRAS-transformed, *p19^Arf^*-deficient MEFs (Fig. S3B). Thus, UPF1 is involved in suppressing C/EBPβ phosphorylation and DNA binding in transformed cells.

**Figure 3.**
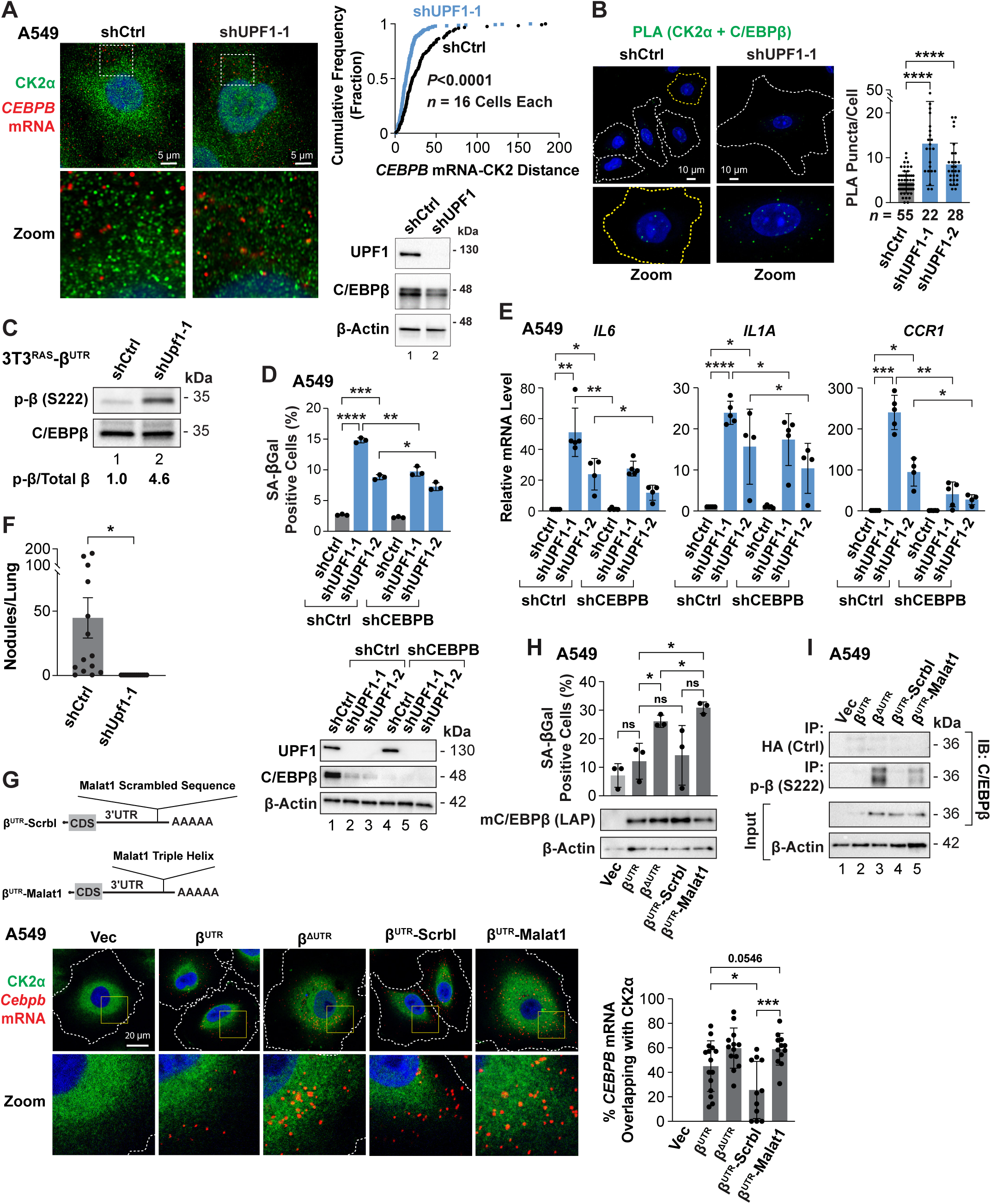
UPF1 depletion in tumor cells stimulates C/EBPβ activity and phosphorylation by CK2α. **(A)** High resolution imaging showing close juxtaposition of *CEBPB* mRNAs and CK2α foci in the perinuclear cytoplasm of UPF1-depleted cells. Right: nearest neighbor analysis of *CEBPB* mRNA and the closest CK2α signal (cumulative frequency plot of distance in pixels). Statistical significance was determined using the Kolmogorov-Smirnov test. **(B)** Proximity ligation assay (PLA) demonstrating increased adjacency of C/EBPβ and CK2α proteins following UPF1 depletion. Right: PLA quantification (PLA puncta per cell). **(C)** Upf1 knockdown increases phosphorylation on C/EBPβ Ser222 (CK2 site). Chromatin bound proteins from 3T3^RAS^-β^UTR^ cells were released by DNase digestion, equalized for total C/EBPβ and analyzed by immunoblotting using a p-Ser222 phospho-specific antibody. **(D)** UPF1 knockdown induces senescence in tumor cells. UPF1 was ablated in A549 cells with or without *CEBPB* knockdown (shCEBPB) and SA-βGal staining was used to analyze senescence (n ≥ 400 cells). **(E)** UPF1 depletion induces expression of SASP genes that is partially C/EBPβ**-**dependent. SASP mRNA levels were analyzed using RT-qPCR. **(F)** 1×10^6^ control or UPF1-depleted A549 cells were injected intravenously into nude mice and lung nodules were counted after seven weeks. *n* = 14 animals per group. **(G)** Insertion of the *Malat1* stability element at the 3’ end of *Cebpb* mRNA disrupts peripheral localization of *Cebpb* transcripts. Mouse *Cebpb* constructs containing the *Malat1* triple helix or a scrambled version inserted at the 3’ end (diagram) were transduced into A549 cells. *Cebpb^UTR^* (βUTR) and *Cebpb^ΔUTR^* (ΔUTR) were included as controls. RNA FISH was performed using a probe specific for mouse *Cebpb*, combined with IF staining for CK2α. Analysis of *Cebpb* mRNA overlap with the CK2 region is shown on the right. **(H)** SA-βGal staining was used to analyze senescence in the cells described in **(G)** (n ≥ 400 cells). **(I)** Analysis of C/EBPβ Ser222 phosphorylation in the cells described in **(G)**. Nuclear lysates were immunoprecipitated with the p-C/EBPβ (S222) antibody followed by immunoblotting for total C/EBPβ. HA antibody was used as an IP isotype control. *n* = number of cells analyzed. Statistical significance (except for the analysis in panel **A**) was determined using Student’s t test. *p < 0.05,**p < 0.01,***p < 0.001,****p < 0.0001.

UPF1 depletion potently induced senescence (SA-βGal positivity) in a partially C/EBPβ-dependent manner in A549 cells (Fig. 3D) and other human tumor cell lines (Fig. S3C). Consistent with the senescent phenotype, pro-inflammatory SASP genes were strongly up-regulated following UPF1 ablation in A549 cells and required C/EBPβ (Figs. 3E and S3D). SASP induction was also seen in PANC1 and SW-900 tumor cells and RAS-transformed MEFs (Fig. S3E). A549 cells lacking UPF1 were unable to form lung lesions after intravenous injection into immune-deficient mice (Fig. 3F). Hence, tumor cells require UPF1 for senescence bypass, a hallmark of cancer, at least partly by inhibiting C/EBPβ activity.

To more directly address whether mRNA decay is critical for subcellular partitioning of *CEBPB* transcripts, we generated mouse *Cebpb* constructs containing the triple helical *Malat1* RNA stability element,^29^ or a scrambled version (Scrbl), inserted at the 3’ end of the 3’UTR (Fig. 3G). These constructs, together with *Cebpb^UTR^* and *Cebpb^ΔUTR^* controls, were expressed in A549 cells. FISH labeling of the transcripts using a probe that detects only murine *Cebpb*, together with CK2α IF, showed perinuclear exclusion of the *Cebpb^UTR^* and *Cebpb^Scrbl^* transcripts (Fig. 3G). However, *Cebpb^ΔUTR^* and *Cebpb^Malat1^*mRNAs displayed perinuclear clustering and increased overlap with CK2α (Fig. 3G, right panel). *Cebpb^ΔUTR^* and *Cebpb^Malat1^* also increased the proportion of senescent cells, whereas *Cebpb^UTR^*and *Cebpb^Scrbl^* maintained a basal level of senescence (Fig. 3H). The senescence phenotype was associated with increased phosphorylation on Ser222 (Fig. 3I). Thus, the *Malat1* RNA stability element can override 3’UTR inhibition, providing further evidence that localized mRNA decay prevents C/EBPβ synthesis in the kinase-rich perinuclear region to suppress its phosphorylation and pro-senescence activity.

### Staufen RNA-binding proteins are essential for *CEBPB* PMD

The double-stranded RBPs, Staufen1 and 2, were also enriched in the GRE-associated fraction (Table S1; Fig. 4A). STAU proteins control mRNA decay and localization,^30^ and STAU-mediated mRNA decay (SMD) requires UPF1.^31^ STAU2 depletion is also synthetically lethal with mutant *KRAS* in colorectal cancer cells.^27^ Therefore, we sought to investigate their possible connection to *CEBPB* PMD. nRIP binding assays showed that Stau1 and 2 associate with *Cebpb* transcripts in NIH 3T3 cells (Fig. 4B). IF/RNA FISH imaging of A549 cells revealed that STAU1 and 2 are concentrated in perinuclear regions that contain UPF1, correlating with exclusion of *CEBPB* transcripts (Fig. 4C). These results suggest that STAU1/2 could be involved in perinuclear *CEBPB* mRNA decay.

**Figure 4.**
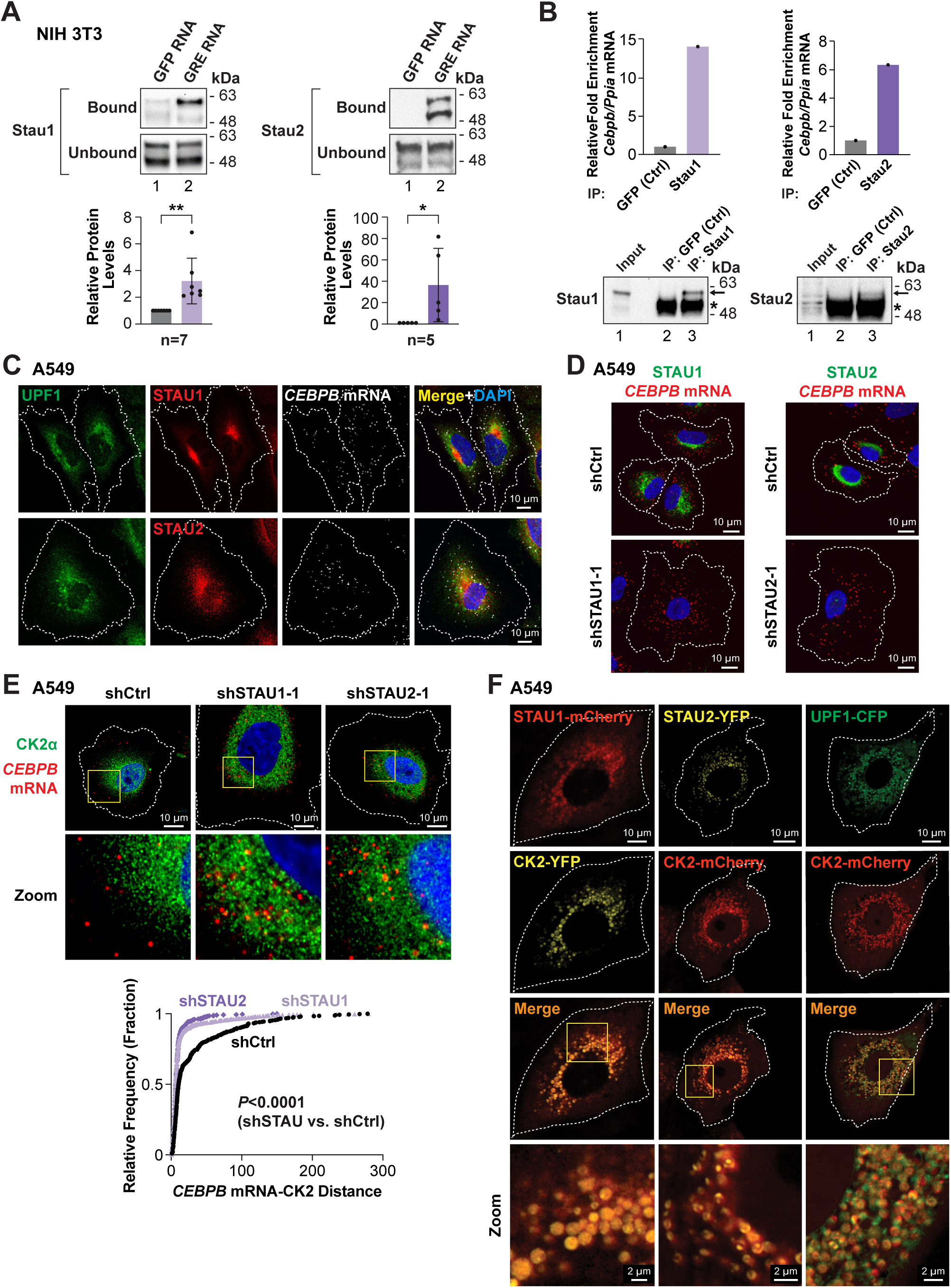
Staufen controls perinuclear exclusion of *CEBPB* mRNA in tumor cells. **A)** Stau1 and Stau2 are enriched in the GRE-bound fraction. Samples described in Fig. 1E were analyzed by immunoblotting for Stau1 or Stau2. **(B)** Stau1/2 associate with *Cebpb* transcripts in NIH 3T3 cells. nRIP assays were performed using Stau1 and Stau2 antibodies and analyzed by RT-qPCR to assess *Cebpb* mRNA enrichment. Bottom: immunoblots for Stau1 and Stau2. **(C)** STAU1 and STAU2 partially overlap with UPF1 in the perinuclear region that lacks *CEBPB* mRNAs. IF staining was combined with *CEBPB* RNA FISH**. (D)** Depletion of STAU proteins disrupts perinuclear exclusion of *CEBPB* mRNAs. Cells were immunostained with the appropriate antibodies to verify STAU knockdown. (**E**) High resolution imaging shows close apposition of *CEBPB* mRNAs and perinuclear CK2α foci in A549 cells upon STAU1 or STAU2 depletion. Bottom: nearest neighbor analysis of CK2α and *CEBPB* mRNAs was performed as described in Fig. 3A. **(F)** Live cell imaging of fluorescently tagged proteins demonstrating overlap of STAU1, STAU2 and UPF1 with perinuclear CK2α signaling endosomes. *n* = number of cells or samples analyzed. Statistical significance (except for the analysis in panel **E**) was determined using Student’s t test. *p < 0.05, **p < 0.01.

Depletion of STAU1 or 2 in A549 cells using shRNAs specific for each isoform (Fig. S4A) increased the perinuclear compartmentalization of *CEBPB* transcripts (Figs. 4D and S4B). High-resolution imaging revealed the presence of *CEBPB* mRNAs closely adjacent to CK2α foci upon STAU1/2 silencing (Fig. 4E), similar to UPF1-depleted cells (Fig. 3A). This finding indicates that STAU1/2 together with UPF1 trigger *CEBPB* mRNA decay on or near CK2 endosomes. Live cell imaging of fluorescently tagged proteins demonstrated that STAU1 and STAU2 form perinuclear foci that almost completely co-localize with co-expressed CK2α on signaling endosomes (Figs. 4F, S4E and S4F). UPF1 also largely overlaps with CK2α, although some UPF1 foci do not coincide with CK2α signals (Fig. 4F). Ablation of either STAU isoform triggered senescence that was partly dependent on C/EBPβ (Fig. S4E) and also induced pro-inflammatory SASP genes (Fig. S4F), although the increase was muted compared to that observed with UPF1 depletion (Figs. 3 and S3). These results support the notion that UPF1 and STAU1/2 regulate *CEBPB* mRNA decay on or near CK2 signaling endosomes to suppress C/EBPβ activation in cancer cells.

### Identification of a STAU binding site (SBS) adjacent to the GRE region

A comparison of human and mouse *CEBPB* 3’UTR sequences shows that the 3’ region encompassing the GRE displays highest conservation (Fig. S5A). An RNA secondary structure model (RNAFold) ^32^ predicts a stem-loop feature formed by part of the GRE paired with a 5’ adjacent sequence (Figs. 5A and S5A,B). A segment of this putative double-stranded region is nearly identical in the two homologs and shows similarity to a known STAU binding site (SBS) in the *ARF1* 3’UTR ^31^ (Fig. S5C). This potential SBS could recruit STAU proteins to the *CEBPB* 3’UTR to activate mRNA decay.

**Figure 5.**
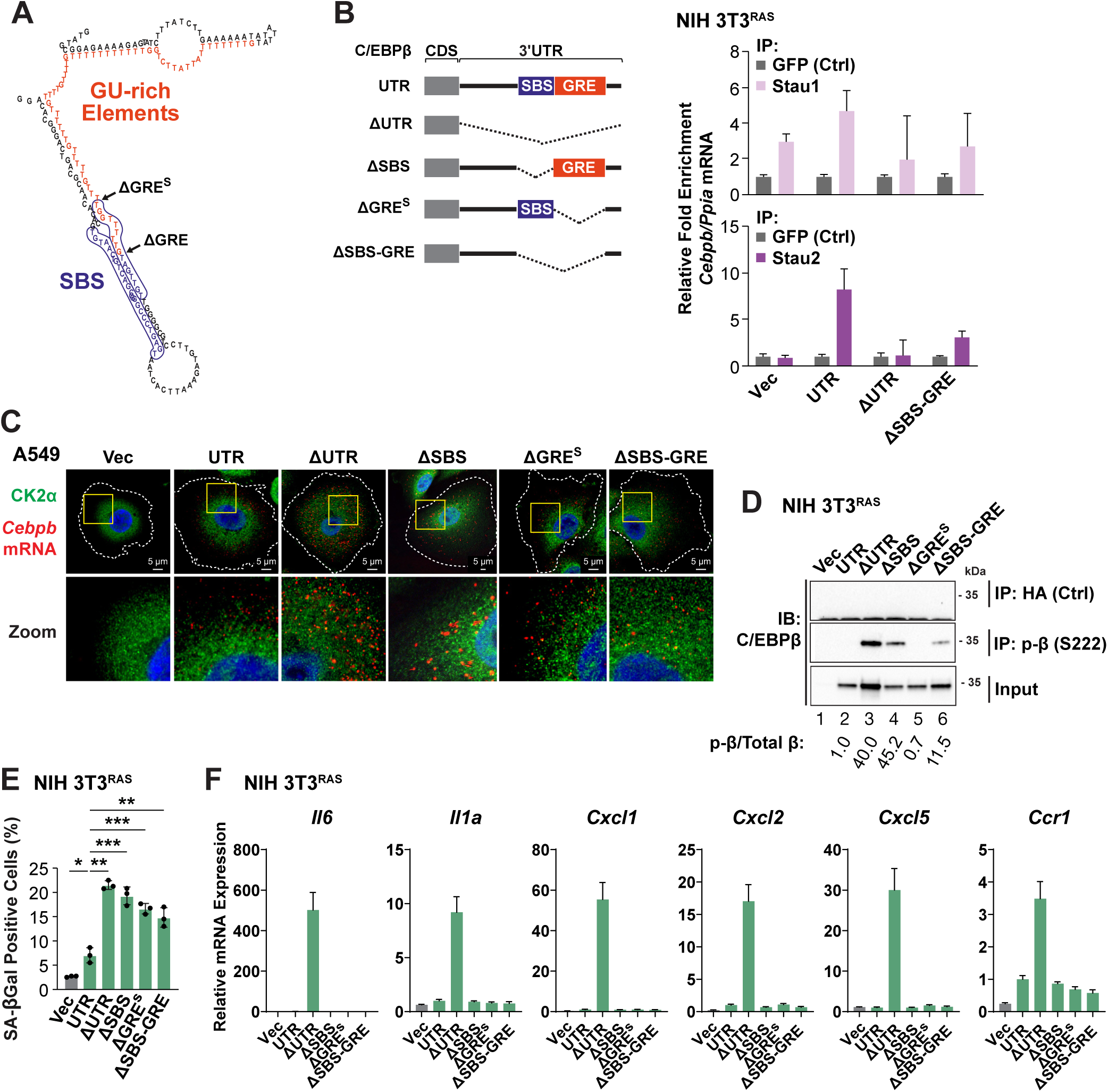
Staufen binds to a putative SBS in the *Cebpb* 3’UTR that is required to suppress C/EBPβ phosphorylation by CK2. **(A)** An RNA secondary structure model (RNAFold) of the murine *Cebpb* 3’UTR predicts a stem-loop feature formed by part of the GU-rich sequence (red) paired with a 5’ adjacent sequence, forming a putative SBS (purple). **(B)** Stau2 binds to the SBS/GRE region in 3T3^RAS^-β^UTR^ cells. Left: diagram of βUTR, ΔUTR, ΔSBS, ΔGRE^S^, and ΔSBS-GRE 3’UTR mutants. Right: nRIP pull-downs were performed using Stau1 and 2 antibodies and analyzed by RT-qPCR to assess *Cebpb* mRNA enrichment for the indicated constructs, normalized to a control mRNA (*Ppia*). **(C)** Analysis of mutants expressed in A549 cells using *Cebpb* mRNA FISH/CK2α imaging. **(D)** *Cebpb* constructs were expressed in 3T3^RAS^ cells and analyzed for C/EBPβ phosphorylation on Ser222. Nuclear lysates were immunoprecipitated using the p-Ser222 antibody followed by immunoblotting for total C/EBPβ. **(E-F)** The same cells were analyzed for senescence (SA-βGal) **(E)** and expression of SASP genes **(F)**.

We generated deletion mutants in murine *Cebpb* that lack the SBS, SBS+GRE and GRE sequences (Fig. 5C). The 5’ endpoint of the latter maintains the full SBS-like sequence and is denoted ΔGRE^S^ to distinguish it from the slightly longer ΔGRE (ARE) deletion described previously ^13^ (Fig. 5A), and which corresponds to our mouse ΔGRE allele (see below). A set of these mutants, together with *Cebpb^UTR^*, were expressed in 3T3^RAS^ cells and nRIP assays performed to assess STAU binding. Stau2 associated with *Cebpb^UTR^* transcripts but showed reduced binding to *Cebpb^ΔSBS-GRE^* and *Cebpb^ΔUTR^* (Fig. 5B). Stau1 binding was less evident and was similar throughout the constructs. Thus, Stau2 binding in particular displays apparent specificity for the SBS element. When expressed in A549 cells, *Cebpb^ΔUTR^* mRNAs overlapped strongly with CK2α as compared with *Cebpb^UTR^* (Fig. 5C). Those lacking the SBS and or GRE also displayed increased localization in the CK2 region. Moreover, when co-transfected with HRAS^G12V^ in HEK 293T cells, *Cebpb^ΔUTR^* and *Cebpb^ΔSBS^* showed strongest phosphorylation on the C/EBPβ CK2 site, Ser222 (Fig. S5D). When expressed in 3T3^RAS^ cells, p-Ser222 levels were increased in mutants lacking the SBS but not the GRE alone (Fig. 5D). However, all except *Cebpb^UTR^* efficiently induced senescence (Fig. 5E) and caused decreased focus formation (Fig. S5E). Strikingly, only *Cebpb^ΔUTR^* stimulated expression of SASP genes in 3T3^RAS^ cells (Fig. 5F). Thus, in transformed cells the SBS suppresses C/EBPβ phosphorylation by CK2, the SBS and GRE motifs inhibit senescence, while a distinct 3’UTR sequence appears to restrain C/EBPβ-induced transcription of SASP genes.

### Oncogenic RAS induces senescence but not the SASP in *p19^Arf−/−^*;*Cebpb-GRE^Δ/Δ^* MEFS

To investigate the *in vivo* relevance of the UPA mechanism, we generated a mouse strain carrying a deletion of the *Cebpb* GRE region, which also removes part of the SBS (Fig. S6A). Homozygous mutant animals (*GRE^Δ/Δ^*) were viable, fertile, and displayed no overt phenotypes. An aging study of *WT* and *GRE^Δ/Δ^* mice revealed no apparent differences in lifespan or susceptibility to disease (Fig. S6B).

To begin to address whether the *GRE^Δ/Δ^* genotype confers resistance to RAS-induced transformation, we crossed the mutant strain to mice lacking the *p19^Arf^*tumor suppressor and prepared MEFs of four genotypes: *WT*, *GRE^Δ/Δ^*, *p19^Arf−/−^*, and *p19^Arf−/−^;GRE^Δ/Δ^*. As C/EBPβ UPA is abrogated in cells undergoing RAS-induced senescence (RIS) but not in RAS-transformed tumor cells,^17^ and *p19^Arf^* deficiency circumvents senescence to facilitate neoplastic transformation, we reasoned that *ΔGRE* phenotypes may be more apparent in cells lacking *p19^Arf^*. RAS-induced phosphorylation on C/EBPβ Ser222 was enhanced ∼2-fold in *p19^Arf−/−^;GRE^Δ/Δ^* MEFs compared to *p19^Arf−/−^* cells (Fig. 6A), consistent with partial disruption of the SBS motif in this allele. *p19^Arf−/−^;GRE^Δ/Δ^* cells also formed fewer RAS-transformed foci than *p19^Arf−/−^* MEFs despite equivalent C/EBPβ levels (Fig. 6B,C). *GRE^Δ/Δ^* MEFS without RAS displayed more SA-βGal positivity than *WT* cells and thus are prone to premature senescence (Figs. 6C and S6C). Upon HRAS^G12V^ expression, senescence of *GRE^Δ/Δ^* cells increased further and was elevated compared to *WT* RIS cells. Whereas *p19^Arf−/−^* cells did not undergo appreciable RIS, *p19^Arf−/−^;GRE^Δ/Δ^* MEFs became senescent and displayed an enlarged, flattened morphology (Fig. S6C). Thus, deletion of the GRE unleashes the RAS-induced pro-senescence activity of C/EBPβ in *p19^Arf−/−^*MEFs.

**Figure 6.**
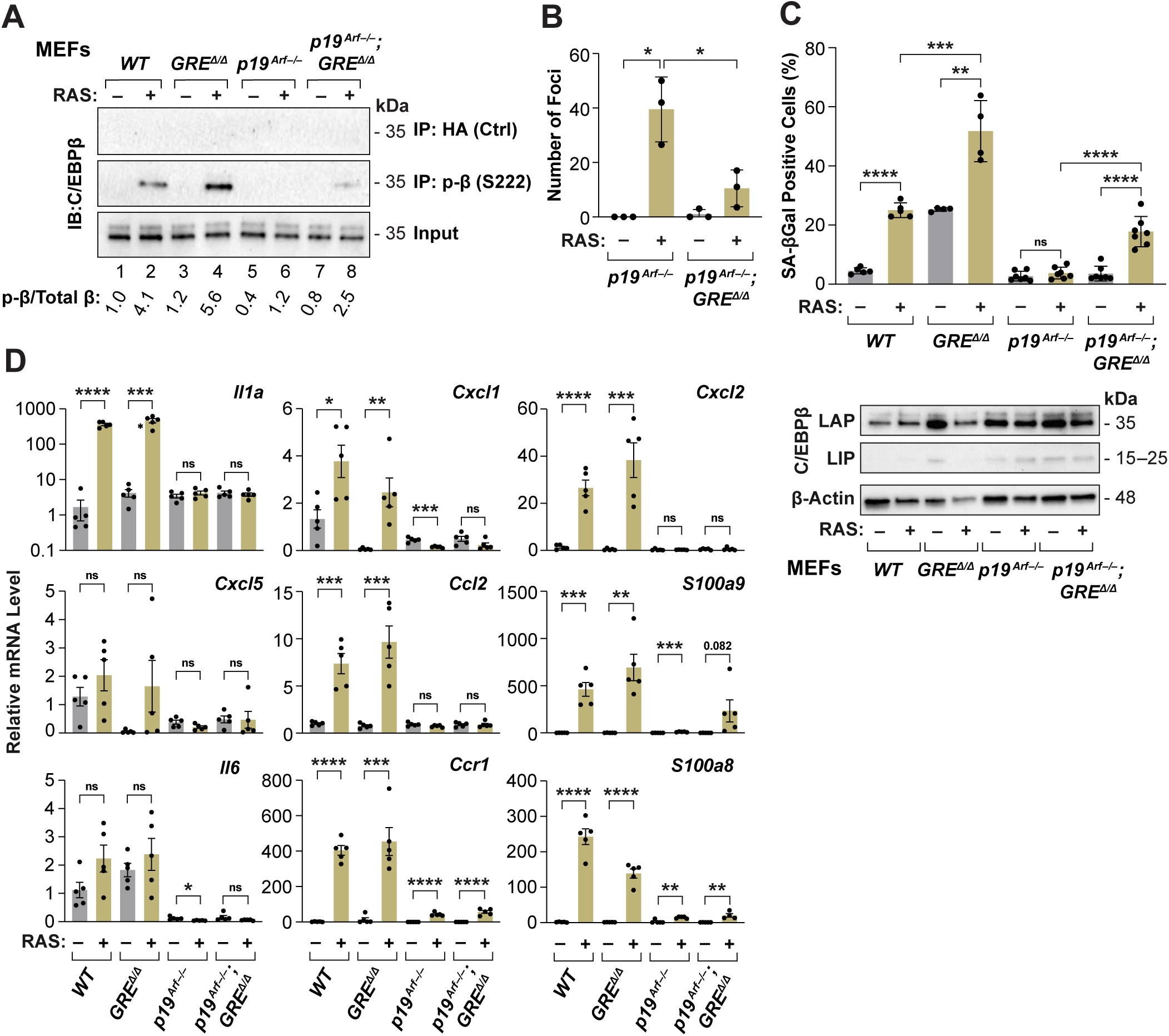
*p19^Arf-/-^*;*Cebpb-GRE^Δ/Δ^* MEFs undergo RAS-induced senescence rather than transformation but do not express the SASP. **(A)** HRAS^G12V^-induced phosphorylation on C/EBPβ Ser222 is enhanced in *p19^Arf-/-^*;*GRE^Δ/Δ^* MEFs. Nuclear lysates from *WT*, *GRE^Δ/Δ^*, *p19^Arf-/-^* and *p19^Arf-/-^*;*GRE^Δ/Δ^* MEFs (± HRAS^G12V^) were analyzed as described in Fig. 5D. **(B)** Focus assays testing neoplastic transformation of *p19^Arf-/-^* and *p19^Arf-/-^*; *GRE^Δ/Δ^* MEFs (± HRAS^G12V^). *WT* and *GRE^Δ/Δ^* cells undergo OIS and do not form transformed foci. Three independent MEF populations for each genotype were analyzed. **(C**) Senescence assays for each MEF population ± HRAS^G12V^. Senescence is the percentage of SA-βGal positive cells (n>400 cells scored) in 4-7 independent MEF cultures. Bottom: immunoblot showing C/EBPβ levels in each MEF population. **(D)** HRAS^G12V^-expressing *p19^Arf-/-^*;*GRE^Δ/Δ^* MEFs only weakly activate canonical proinflammatory SASP genes but induce *S100a9*. RNA samples from *WT*, *GRE^Δ/Δ^*, *p19^Arf-/-^*, and *p19^Arf-/-^*;*GRE^Δ/Δ^* MEFs (± HRAS^G12V^) were analyzed by RT-qPCR for a panel of SASP genes. All values were normalized to the average of the *WT* controls (without HRAS^G12V^). Statistical significance was determined using Student’s t test. *p < 0.05,**p < 0.01,***p < 0.001,****p < 0.0001; ns, not significant.

HRAS^G12V^ stimulated expression of pro-inflammatory SASP genes equivalently in *WT* and *GRE^Δ/Δ^* MEFS but not in *p19^Arf−/−^* cells (Fig. 6D). Notably, *p19^Arf−/−^;GRE^Δ/Δ^* MEFs also failed to express the SASP despite becoming senescent. This phenotype is similar to that of 3T3^RAS^ cells over-expressing *Cebpb* 3’UTR mutants lacking the GRE and/or SBS elements (Figs. 5E,F) and provides further evidence that an independent 3’UTR sequence inhibits the ability of C/EBPβ to activate transcription of SASP genes. A notable exception was S100a9, which was induced by RAS in *p19^Arf−/−^;GRE^Δ/Δ^* MEFs but only minimally in *p19^Arf−/−^* cells (Fig. 6D). S100a9 is a secreted Ca^++^ binding protein with pro-inflammatory properties that binds to TLR4 and activates the inflammasome.^33^ S100a9 also induces senescence in bone marrow mesenchymal cells^34^ and could possibly act as an autocrine/paracrine RIS effector in *p19^Arf−/−^;GRE^Δ/Δ^* cells.

### *Kras^G12D^*-driven lung tumors in *Cebpb-GRE^Δ/Δ^* mice show impeded progression to adenocarcinoma

To extend our findings to an *in vivo* setting, we crossed *GRE^Δ^* mice to the cancer prone strain *KRas^LA2^*, which develops lung tumors with high penetrance.^35^ Pulmonary lesions in this model are predominantly benign adenomatous hyperplasias and alveolar adenomas, although some of these progress to malignant adenocarcinomas (ADCs). Groups of *KRas^LA2^/+* and *KRas^LA2^/+;GRE^Δ/Δ^* mice were aged until pulmonary tumor burdens or other lesions required euthanasia. There was no difference in survival between the two genotypes (Fig. 7A). Overall tumor burdens (sum of all tumor areas) showed a modest reduction in *GRE^Δ/Δ^* mice that was not significant (*p*=0.09) (Fig. 7B). However, histopathological evaluation of lesions showed differences in the most severe grade observed, as ADCs were the most advanced tumors seen in 71.4% of *WT* mice compared to just 14.3% in the *GRE^Δ/Δ^* cohort (*p*=0.0016; Fisher’s exact test) (Fig. 7C). Using a supervised, AI-based algorithm (HALO image analysis software; Indica Labs) trained on H&E images to distinguish benign adenomas from atypical cells/ADCs, we quantitated tumor areas corresponding to each grade (Fig. 7C). Although adenomas were similar between the two genotypes, areas of adenocarcinoma were nearly 4-fold greater in *WT* mice than *GRE^Δ/Δ^* animals (*p*=0.0244). This difference was not attributable to age at sacrifice, as there was no correlation between age and ADC burden in either genotype (Fig. S7A).

**Figure 7.**
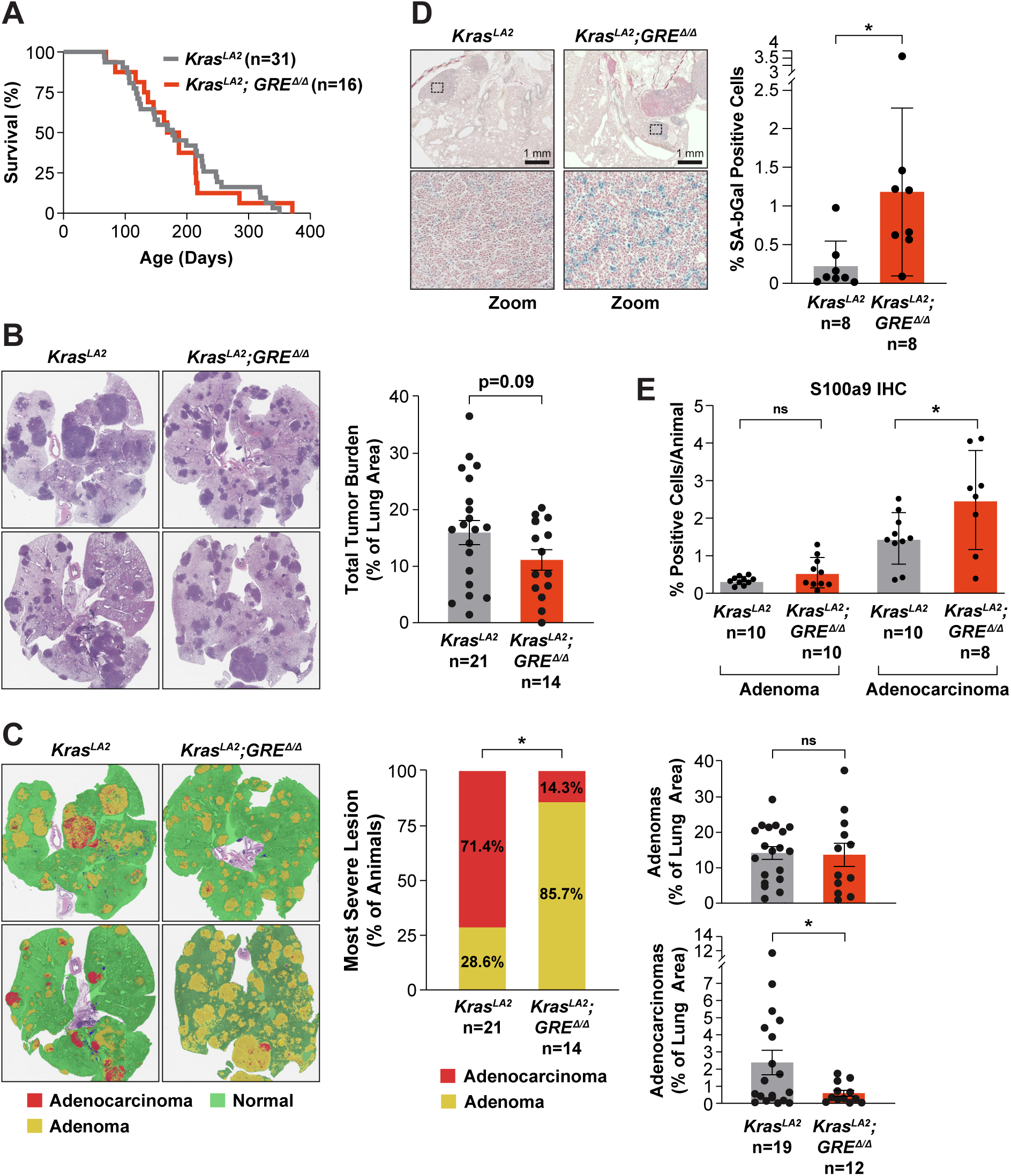
*Cebpb-GRE^Δ/Δ^* mice display impaired progression to adenocarcinoma in a *Kras^G12D^*-driven model of lung cancer. **(A)** Kaplan-Meier curves of *Kras^LA2/+^*and *Kras^LA2/+^;GRE^Δ/Δ^*mice. No significant difference in survival was observed (Log-rank test; p = 0.84). **(B)** Left: two representatives H&E images of tumor-bearing lungs from *Kras^LA2/+^*and *Kras^LA2/+^;GRE^Δ/Δ^* mice. Right: total lung tumor burdens (percent tumor area/animal) at clinical sacrifice. Tumor areas were determined using the HALO random forest tissue classifier. **(C)** Left: two representative HALO AI segmentations overlaid on H&E images from *WT* and *GRE^Δ/Δ^* lungs (same specimens shown in panel B). HALO AI identifies and segments normal lung tissue (green), adenomas (yellow) and adenocarcinomas (red). Middle: Most severe tumor grades in *Kras^LA2/+^*and *Kras^LA2/+^*;*GRE^Δ/Δ^* mice as determined by pathological examination of lesions. Data are shown as the percentage of animals in each group (Fisher’s exact test; p = 0.0016). Right: HALO AI quantification of adenoma (top) and adenocarcinoma (bottom) burdens (percent lesion area per animal). **(D)** Senescent cells in *Kras^LA2^* and *Kras ^LA2^;GRE^Δ/Δ^* tumors. Lung sections were stained for SA-βGal (left) and scanned to quantitate positive cells (right). Data are expressed as % positive cells in tumor areas per animal. **(E)** S100a9 positive cells in *Kras^LA2^* and *Kras ^LA2^;GRE^Δ/Δ^* tumors. Lung sections were analyzed by IHC staining for S100a9. Data are expressed as % positive cells in tumor areas per animal, stratified by tumor grade. *n* = number of animals analyzed. Statistical significance was determined using Student’s t test, except where otherwise indicated. *p < 0.05; ns, not significant.

Ki67 staining in ADC areas did not differ between the two genotypes (Fig. S7B), indicating that rare tumors that evade the *GRE^Δ/Δ^* blockade are able to achieve a high proliferation rate. There was also no difference in infiltration of M1 and M2 macrophages in WT and *GRE^Δ/Δ^* tumors (Fig. S7C). However, quantitation of SA-βgal positive areas in tumor tissues revealed significantly more senescence in *GRE^Δ/Δ^* lesions (Fig. 7D), providing an explanation for the decrease in ADC burdens in the mutant cohort. *GRE^Δ/Δ^* tumors also contained more S100a9 positive cells (Fig. 7E), consistent with the increased RAS-induced expression of this cytokine in *GRE^Δ/Δ^* MEFs. A characteristic feature of cancer cells is variability in nuclear size and shape. To determine if *WT* and *GRE^Δ/Δ^* tumors exhibit differences in anisokaryosis, adenoma and ADC regions were analyzed for nuclear size heterogeneity using HALO AI. While distributions of nuclear size were similar in adenomas, *GRE^Δ/Δ^* ADC lesions contained significantly fewer karyomelagic cells compared to *WT* tumors (p<0.0001) (Fig. S7D). These findings support the reduced malignancy of *GRE^Δ/Δ^* cancers.

## Discussion

UPA inhibits C/EBPβ activity in tumor cells through 3’UTR-mediated exclusion of *CEBPB* transcripts from the kinase-rich perinuclear region.^13^ We now identify perinuclear mRNA decay (PMD) as a mechanism for eliminating *CEBPB* mRNAs from the nuclear-proximal compartment. UPF1 and STAU, which are well documented mRNA decay proteins, play key roles in PMD. The *CEBPB* GRE sequence and an adjacent SBS motif recruit HuR and STAU, respectively, to control mRNA localization. However, these two elements act differently, as the SBS inhibits C/EBPβ phosphorylation by CK2 whereas the GRE does not.

The subcellular location of PMD is essential to suppress C/EBPβ phosphorylation in tumor cells, as the C/EBPβ kinases CK2 and ERK are present on perinuclear signaling endosomes.^12^ Localized *CEBPB* mRNA decay is determined by perinuclear compartmentalization of UPF1 and STAU, whose presence is inversely correlated with *CEBPB* transcripts. Depletion of UPF1 or STAU1/2 caused many *CEBPB* transcripts to colocalize with CK2 puncta (Figs. 3A and 4F). Live cell imaging demonstrated strong overlap of STAU1 and STAU2 with CK2 puncta, and partial overlap of UPF1 and CK2 (Fig. 4F). This implies that *CEBPB* mRNA decay is highly localized to signaling endosomes. Therefore, PMD may be more accurately described as kinase-proximal mRNA decay (KPMD). The co-incidence of CK2 and STAU provides a compelling explanation for the increased phosphorylation on C/EBPβ Ser222 when the SBS motif is deleted but not when the GRE alone is absent. These findings also indicate that despite their partially overlapping configuration, the SBS does not require the GRE to recruit STAU2 or to initiate mRNA decay.

The presence of *CEBPB* transcripts near CK2 endosomes when KPMD is disrupted suggests the existence of an intrinsic homing mechanism for *CEBPB* mRNAs. Since *CEBPB* transcripts lacking the 3’UTR also localize near CK2, there could be an RNA trafficking determinant within the coding region. Alternatively, a localization signal within the nascent C/EBPβ protein might direct *CEBPB* polysomes to CK2-containing endosomes. The proximity between *CEBPB* mRNAs and CK2 suggests a spatially coupled phosphorylation mechanism, whereby modifications are associated with protein synthesis (Fig. S7C). This coupling would explain why C/EBPβ translated from transcripts located outside the perinuclear domain remains in an unphosphorylated, low activity state. The default propensity of *CEBPB* mRNAs to localize near CK2 signaling endosomes underscores the importance of KPMD in preventing C/EBPβ phosphorylation/activation to suppress senescence in tumor cells.

An unexpected finding of our study was that the pro-senescence and SASP-inducing activities of C/EBPβ are inhibited by distinct 3’UTR elements. This is clearly demonstrated by the different responses of transformed cells to expression of *Cebpb^UTR^* (neither senescence nor the SASP), ΔGRE and/or SBS (senescence only), or *Cebpb^ΔUTR^* (senescence and the SASP). Further evidence comes from the senescence-prone but SASP-less phenotype of immortalized *GRE^Δ/Δ^* MEFs expressing oncogenic RAS. Hence, while the SBS/GRE region is required to inhibit senescence, deletion of this sequence does not lead to activation of SASP genes, implicating another 3’UTR element in suppressing the latter. This conclusion is reinforced by the fact that HuR depletion induces senescence but only weakly activates the SASP (Fig. S1B), similar to deletion of its cognate GRE site. By contrast, UPF1 silencing activates both C/EBPβ-dependent senescence and the SASP, indicating that mRNA decay is central to both inhibitory mechanisms.

*GRE^Δ/Δ^* mice exhibited no obvious phenotypes, indicating that at least the GRE-dependent functions of UPA are not essential for normal mouse development or physiology. However, in a *Kras^G12D^*-driven model of lung carcinogenesis, the *ΔGRE* mutant suppressed development of malignant adenocarcinomas and caused increased senescence in tumor areas, mirroring the results in *p19^Arf−/−^;GRE^Δ/Δ^* MEFs. These findings provide the first evidence that UPA inhibits the tumor suppressor activity of C/EBPβ *in vivo*. Development of benign adenomatous lesions was unaffected, indicating that UPA becomes critical when tumors progress to the carcinoma stage. *Kras^LA2^*-driven lung ADCs are characterized by elevated RAS pathway signaling and increased p-ERK levels compared to adenomas.^36^ Hyperactive RAS activates C/EBPβ, which would impede growth of adenocarcinomas without the constraint of UPA. Amplified RAS signaling also engages pro-senescence pathways such as Arf/p53,^37^ creating selective pressure for loss of tumor suppressor genes in malignant cancers.^36^ We propose that such events, particularly loss of p19*^Arf^*, are linked to implementation of C/EBPβ UPA. This is supported by the observation that the *GRE^Δ/Δ^* genotype restores RAS-induced senescence in *p19^Arf−/−^* MEFs (Fig. 6C). The UPA mechanism could extend to other tumor suppressor proteins, whose activities may be similarly constrained by 3’UTR regulation.

## Supporting information

Supplementary Figures and legends and critical reagents list

## Acknowledgements

We thank Kristen Lynch for insightful comments, Eugene Valkov for suggesting the *Malat1* experiment and providing constructs, Dominic Esposito for RAS vectors, and Allen Kane for preparation of figures. This research was supported in part by the Intramural Research Program of the NIH, National Cancer Institute, Center for Cancer Research, and in part with Federal funds from the National Cancer Institute, National Institutes of Health, under Contract No. HHSN261200800001E. The content of this publication does not necessarily reflect the views or policies of the Department of Health and Human Services, nor does mention of trade names, commercial products, or organizations imply endorsement by the U.S. Government.

## Methods

### Lead contact

Further information and requests for resources and reagents should be directed to and will be fulfilled by the lead contact, Peter F. Johnson.

### Materials availability

Upon request, cell lines, mouse strains and plasmids generated in the lab will be made available.

### Cell lines

Mouse NIH 3T3 and NIH 3T3^RAS^ fibroblasts were cultured in DMEM supplemented with 10% calf serum. Mouse embryonic fibroblasts (MEFs), human A549, PANC-1, SW-900, IMR-90, GP2 and HEK 293T cell lines were cultured in DMEM supplemented with 10% fetal bovine serum. All cell lines were cultured with 100 µg/mL primocyn, maintained at subconfluency and grown in a humidified incubator at 37 °C with 5% CO_2_. Cells infected with retroviral/lentiviral plasmids were selected with 2 µg/mL puromycin for 2 days or 100 µg/mL hygromycin for 4 days or 800 µg/mL G418 for 7 days.

### Animal studies

All animals were maintained in compliance with National Institutes of Health animal guidelines and experimental procedures were approved by the Animal Care and Use Committee, National Cancer Institute at Frederick. *p19^Arf-/-^* (*CdKn2a-Arf*) mice were obtained from the NCI-Frederick repository on a mixed background (B6.129). This strain was continuously backcrossed to C57Bl/6Ncr and maintained as homozygous stocks. *Cebpb-GRE^flox^* mice were generated in house by injection of ES cells containing the targeted *GRE^flox^* allele (*LoxP* sites flanking 102-nt mouse *Cebpb* GRE region; 3’UTR nucleotides 308-409) into C57BL/6Ncr mouse blastocysts. The germline GRE^Δ^ allele was generated by mating *GRE^flox^* mice with *ACTB-Cre* mice ^38^. GRE^Δ^ mice were backcrossed to *WT* C57BL/6Ncr animals and maintained as heterozygous stocks. *p19^Arf-/-^; GRE^Δ/+^* strain (C57BL/6Ncr) was obtained by continued intercrossing of *p19^Arf-/-^ and GRE^Δ/+^* mice and maintained by *p19^Arf-/-^; GRE^Δ/+^* crosses. *Kras^G12D-LA2/+^* mice ^35^ were maintained by crossing to *WT* (129/SV). To obtain the *Kras^LA2/+^;GRE^Δ/+^*strain, *Kras^LA2/+^* (129/SV) and *GRE^Δ/+^* (C57BL/6Ncr) mice were crossed. The mixed B6.129 background F1 progeny were bred (*Kras^LA2/+^;GRE^Δ/+^*X *Kras^+/+^;GRE^Δ/+^*) to produce experimental animals. Mouse embryonic fibroblasts (MEFs) were prepared from embryonic day 13.5 (E13.5) embryos of the indicated genotypes.

### Xenograft experiments

Six- to eight-week-old Balb/c athymic nude mice were randomly grouped. The mice were warmed with an incandescent lamp to induce vasodilation, placed in a commercial restraint device, and injected via the lateral tail vein with 1 × 10^6^ A549 cells or derivatives suspended in 100 µl of phosphate buffer saline (PBS). Animals were euthanized 7 weeks post-injection using CO_2_ inhalation. The entire lung and bronchi were harvested, lung nodules were counted and measured. The lungs were inflated using 10% Neutral Buffered Formalin (NBF) before placing into a container of the same fixative. To ensure proper fixation stability, after five days in fixative, the tissues were removed from 10% NBF and placed into 70% ethanol.

### Plasmids

Oligonucleotides encoding short hairpin transcripts targeting mouse *Anxa1, Anxa2, Celf1, Dhx9, Fmr1, Fubp1, Fubp2, Fubp3, Hnrnpa2b1, Hnrnpd, Igf2bp1, Igf2bp2, Igf2bp3, Ilf2, Ilf3, Mapre1, Nono, Plec, Strbp* and *Upf1* (Table S1) were inserted between the BamHI and EcoRI sites of the knockdown vector pSirenRetroQ-puro (Clontech). The knockdown vectors for mouse *Cebpb* (Cebpb1 shRNA) and human *CEBPB* have been described.^22^ The expression vectors pcDNA3.1(-)-β^UTR^, pcDNA3.1(-)-β^ΔUTR^, pBabe-β^UTR^, pBabe-β^ΔUTR^, and knockdown vector pSuperRetro-shElavl have been described previously.^13^ Lentiviral plasmids CMV51p-HRAS^G12V^ (hygro) and control CMV13p (hygro) were obtained from Dr. Dominic Esposito (RAS Initiative, Frederick National Laboratory for Cancer Research). The triple helix sequence from mouse noncoding RNA *Malat1* ^29^ and its control scrambled sequence were inserted into the 3’ end of pBabe-βUTR using synthetic oligos harboring PacI and SbfI restriction sites. The ΔSBS, ΔGRE^S^, ΔSBS-GRE deletions were generated by replacing the sequence listed in Table S1 with a BamHI site. Briefly, two PCR products, containing NheI/BamHI 5’ fragment and BamHI/HindIII 3’ fragment, were cloned into pcDNA3.1(-)-β^ΔUTR^ by three-way ligation. Each sequence was transferred from pcDNA to pBabe retroviral vector using EcoRI and SalI sites. A complete list of plasmids and sequences is included in Table S1.

### Retroviral and lentiviral infections

Packaging GP2 cells (for retroviral plasmids) or HEK 293T cells (for lentiviral plasmids) were seeded at 3 × 10^6^ cells per 100 mm culture dish for 24 h before transfection with 20 µg of the viral expression plasmid containing the insert of interest. Plasmids were co-transfected by calcium phosphate precipitation together with the following packaging and/or envelope plasmids: 4 µg of pVSVG (for retroviral plasmids) or 6 µg of pMD2.G, 10 µg of pMDL g/p RRE, 5 µg of pRSV-Rev (for lentiviral plasmids). Supernatants containing infectious viral particles were harvested at 24, 36 and 48 h post-transfection and filtered through 0.45 µm PVDF filters. Cells were cultured in 100 mm culture dishes for 24 h before infection with viral supernatants in the presence of 8 µg/mL polybrene.

### Transient transfection

HEK 293T cells were transfected with 100 ng of pcDNA 3.1(-)-HRAS^G12V^, 1 µg of pcDNA 3.1(-) empty vector or C/EBPβ deletion mutants in 100 mm dishes using X-tremeGENE HP (Roche) according to manufacturer’s instructions. After transfection, cells were cultured for 48 hours before harvesting.

### Protein lysates

For cytoplasmic and nuclear fractionation, cells were rinsed twice with ice-cold PBS, resuspended in hypotonic Buffer A (20 mM HEPES [pH 7.9], 1 mM EDTA, 10 mM NaCl, 1 mM DTT, 0.1% NP-40 substitute, 1X protease and phosphatase inhibitor sets) and incubated on ice for 10 min. Nuclei were pelleted by centrifugation for 10 min at 1,200 x g at 4 °C. Supernatants (cytoplasmic extract) were frozen at −80 °C. Nuclear pellets were rinsed with Buffer A (without NP-40) to remove residual cytoplasmic contaminants. Nuclear proteins were extracted by incubation of nuclei in hypertonic Buffer C (25 mM HEPES [pH 7.9], 0.2 mM EDTA, 700 mM NaCl, 0.2 mM DTT, 25% glycerol, 1X protease and phosphatase inhibitor sets) for 30 min at 1,300 rpm at 4 °C. Nuclear debris was pelleted by centrifugation for 10 min at 12,000 x g at 4 °C, and the supernatant (nuclear extract) was stored at −80 °C. Chromatin fractionation was performed as described with minor modifications.^39,40^ Briefly, cells were rinsed twice with ice-cold PBS, resuspended in Buffer A* (10 mM HEPES [pH 7.9], 10 mM KCl, 1.5 mM MgCl_2_, 340 mM sucrose, 10% glycerol, 0.1% NP-40, 1 mM DTT, 1X protease and phosphatase inhibitor sets) and incubated on ice for 10 min. Nuclei were pelleted by centrifugation for 5 min at 1,200 x g at 4 °C. Obtained supernatant was labeled as cytoplasmic extract and frozen at −80 °C. Nuclear pellets were rinsed with Buffer A* (without NP-40) to remove residual cytoplasmic contaminants. Nuclear pellet was incubated with Buffer B (3 mM EDTA, 0.2 mM EGTA, 1 mM DTT, 1X protease and phosphatase inhibitor sets) for 30 min at 1,300 rpm (thermomixer) at 4 °C. Insoluble chromatin fraction was collected by centrifugation for 4 min at 1,300 x g at 4 °C, and the supernatant was designated as nucleoplasmic extract and stored at −80 °C. Insoluble chromatin fraction was gently rinsed three times with ice-cold Buffer B and collected by centrifugation for 4 min at 1,300 x g at 4 °C. Chromatin-bound proteins were obtained by incubation of insoluble chromatin fraction with DNase I solution (0.1 U/μL DNase I, 10 mM Tris-HCl pH 7.5, 2.5mM MgCl_2_, 0.1 mM CaCl_2_, 1X protease and phosphatase inhibitor sets) at 37 °C for 1 h. Soluble chromatin fraction was cleared by centrifugation at 16,000 × g, 10 min, 4 °C and supernatant was frozen at −80 °C. For whole cell lysates, cells were rinsed twice with ice-cold PBS, harvested in RIPA buffer (10 mM Tris pH 7.4, 150 mM NaCl, 1 mM EDTA, 0.5% sodium deoxycholate, 0.1% SDS, 1% Triton-X, 1X protease and phosphatase inhibitors sets), kept on ice for 10 min, and centrifuged for 10 min at 12,000 x g at 4 °C. Supernatant was stored at −80 °C. Protein concentrations were determined by Bradford protein assay (Bio-Rad).

### RNA affinity purification of GRE interacting proteins

The RNA pull-down method was modified from Butter et al.^20^ The GRE and GFP sequences were fused to the S1 aptamer tag, which was selected to have high affinity for streptavidin.^41^ The GRE and GFP templates for *in vitro* transcription (TranscriptAid T7 High Yield Transcription Kit, ThermoFisher) were purified PCR products made with oligonucleotides containing the T7 RNA polymerase promoter sequence at the 5’ end and the aptameric tag at the 3’ end (Table S1). The RNA was purified using phenol/chloroform standard protocols and stored in RNase-free water at −80 °C. The S1-tagged RNA was refolded by heating for 3 min at 95 °C, cooling on ice for 2 min, adding the RNA buffer (10 mM Tris-HCl [pH 8.0], 150 mM NaCl, 1 mM MgCl_2_, 0.1 mM EDTA) and incubating for 20 min at 20 °C. The RNA was bound to RNase-free Dynabeads MyOne Streptavidin C1 (ThermoFisher) in RNA binding buffer (RBB, 50 mM Hepes·[pH 7.4], 150 mM NaCl, 0.5% NP-40 substitute, 10 mM MgCl_2_) by incubation with gentle rotation for 30 min at 4 °C, followed by three washes with RBB. Beads were incubated with 400 μg NIH 3T3 cytoplasmic extract (in Buffer A with NaCl concentration adjusted to 150 mM and prepared with RNase inhibitor), 40 U of RNase inhibitor (ThermoFisher), and 20 μg of yeast tRNA (ThermoFisher) for 30 min at 4 °C (final volume 100 μL). After three RBB washes with gentle rotation for 10 min each at 4 °C, the RNA was competed from the beads with elution buffer (16 mM biotin, 50 mM Hepes [pH 7.4], 250 mM NaCl, 0.5% NP-40 substitute, 10 mM MgCl_2_). Eluates were frozen at −80 °C until used for immunoblotting or MS analysis.

### Mass spectrometry (LC-MS/MS) to identify GRE interacting proteins

Coomassie stained gel bands were chopped to small pieces and destained using 50% CAN, 25 mM NH_4_HCO_3_ [pH 8.4]. After removal of organic solvent, the gel pieces were vacuum dried for 45 min. The dried gel pieces were rehydrated in 50 µL of trypsin (20 ng/ µL) resuspended in 25 mM NH_4_HCO_3_ and incubated on ice for 30 min. Excess trypsin was removed and replaced with 30 μL of 25 mM NH_4_HCO_3_ [pH 8.4]. The samples were incubated at 37 °C for 16-18 hr. Peptides were extracted from the gel bands with 70% (v/v) CAN, 0.1% TFA. The extracted peptides were lyophilized, desalted using C18 columns (ThermoFisher) and reconstituted in 0.1% TFA. The peptides were analyzed using mass spectrometry as described below. The samples were subjected to nanoflow liquid chromatography on a 25 cm Acclaim PepMap C18 column (Easy nLC 1200, Thermo Scientific) coupled to high resolution tandem mass spectrometer (Q Exactive HF, ThermoFisher). MS precursor scans were performed at a resolution of 60,000 with an ion accumulation target set at 1e^6^, maximum IT at 120 ms, over a mass range of 380-1580 m/z. MS2 analysis was performed at a resolution of 15,000 with an ion accumulation target set at 2e^5^, maximum injection time at 50 ms, at an isolation with of 1.4 m/z over a scan range of 200 to 2000 m/z. Top 20 precursor ions were selected for fragmentation at a normalized collision energy of 27, charge state 1 and >8 were excluded with a 25 second dynamic exclusion. Acquired MS/MS spectra were searched against the mouse Uniprot protein database using SEQUEST and the Percolator validator algorithms in Proteome Discoverer 2.2 (ThermoFisher). The precursor ion tolerance was set at 10 ppm and the fragment ions tolerance at 0.02 Da. Maximum 2 missed cleavage side were allowed, a minimum peptide length was set at 6 amino acids with methionine oxidation as dynamic modification. A false discovery rate of 0.01 was used for the decoy database search for peptide identification.

*MS data analysis.* The MS results identified 25,188 PSMs bound to GRE representing 1107 proteins, and 13,352 PSMs bound to GFP representing 763 proteins (the sum of PSM counts from three independent affinity purification experiments was used). Overall, there were 4616 identified proteins that bind to either GPF or GRE. Since the GRE and GFP pull down experiments were run independently, the null hypothesis is that there should be no preference for binding to one or the other. By determining the number of PSMs from a protein that bound to GRE and the total number that were bound to either GRE or GFP, a Binomial test was used to determine if the number of GRE-bound PSMs is significantly different from that expected by chance. This p-value was Bonferroni corrected and then used to calculate the false discovery rate (FDR). Using a threshold FDR of 0.1, all proteins whose ratio of GRE-bound PSMs to GFP-bound PSMs was at least 2.0 were selected. The 324 proteins that show a significant preference for binding to GRE were submitted to DAVID (2021 Release)^21^ for functional analyses.

### Immunoblotting

Equal amounts of total proteins were separated on 4-15% SDS-PAGE (Bio-Rad) and blotted to PVDF membranes (Bio-Rad). Membranes were blocked for 1 h in TBS-T buffer (20 mM Tris [pH 7.6], 150 mM NaCl, 0.1% Tween-20) containing 5% nonfat milk and were then incubated with primary antibodies for 16-18 h at 4 °C. Membranes were rinsed three times in TBS-T, incubated for 1 h at 20 °C with the appropriate secondary antibodies in 5% nonfat milk (in TBS-T), then rinsed three times in TBS-T before imaging using chemiluminescent ECL substrate (ThermoFisher). All antibodies are listed in Table S1.

### Immunoprecipitation (IP)

400 µg of cytoplasmic extract (where noted, with 200 µg/mL RNase A treatment for 10 min at 20 °C) or 200 µg nuclear proteins were incubated with primary antibodies with gentle rotation for 3 h at 4 °C in IP buffer (20 mM Tris [pH 8.0], 137 mM NaCl, 10% glycerol and 1% NP-40 substitute, 1X protease and phosphatase sets). Dynabeads™ Protein G (ThermoFisher) were added to the samples and incubated with gentle rotation for 1 h at 4 °C. Immunocomplexes were washed four times in IP wash buffer (20 mM Tris [pH 8.1]; 137 mM NaCl; 0.5% NP-40; 1 mM EDTA) and eluted in 2X Laemmli sample buffer (Bio-Rad) with β-mercaptoethanol for 10 min at 95 °C. Eluted samples were analyzed by immunoblotting. In the case of STAU proteins, immunocomplexes were eluted in 2X Laemmli sample buffer (Bio-Rad) with β-mercaptoethanol for 30 min at 20 °C and Protein A conjugated with HRP (MilliporeSigma) was used in lieu of secondary antibody.^42^ Antibodies are listed in Table S1.

### Native RNA immunoprecipitation (nRIP)

400 µg of cytoplasmic extracts (in Buffer A with 150 mM NaCl, 1 mM MgCl_2_ and prepared with 400 U/mL RNase inhibitor) were incubated with primary antibodies (Table S1) with gentle rotation for 3 h at 4 °C in NT-2 buffer (50 mM Tris [pH 7.4], 150 mM NaCl, 1 mM MgCl_2_, 0.05% NP-40 substitute, 400 U/mL RNase inhibitor, 1X protease and phosphatase sets). Dynabeads™ Protein G (ThermoFisher) were added to the samples and incubated with gentle rotation for 1 h at 4 °C. Immunocomplexes were washed four times in NT-2 buffer, processed for RNA extraction/purification, and analyzed by RT-qPCR with sequence-specific primers (Table S1). For confirmation of immunoprecipitated proteins, the same protocol was performed in parallel except that after washes, beads were eluted in 2X Laemmli sample buffer (Bio-Rad) with β-mercaptoethanol for 10 min at 95 °C. Eluted samples were analyzed by immunoblotting.

### Crosslinking RNA immunoprecipitation (CLIP)

The CLIP procedure was modified from Niranjanakumari et al.^43^ and Yoon and Gorospe.^44^ For STAU1 CLIP, cells were rinsed twice in PBS and 0.7% formaldehyde (in PBS) was added directly to the dishes. Cells were incubated for 10 min at 20 °C with rocking and the cross-linking reaction was quenched with 125 mM glycine for 5 min at 20 °C. Cells were extensively washed with ice-cold PBS, resuspended in ice-cold NT-2 Buffer (50 mM Tris [pH 7.5], 150 mM NaCl, 1 mM MgCl_2_, 0.05 % NP-40, 1X protease and phosphatase inhibitor sets) and kept for 10 min on ice. Lysates were treated with 200 U/mL MNase in the presence of 5 mM CaCl_2_ for 5 min at 37 °C to obtain RNA fragments. After incubation, 10 mM EGTA was added to inactivate MNase reaction. For STAU2 CLIP, cells were rinsed twice in PBS and 3.7% formaldehyde (in PBS) was added directly to the dishes. Cells were incubated for 10 min at 20 °C with rocking and the cross-linking reaction was quenched with 125 mM glycine for 5 min at 20 °C. Cells were extensively washed with ice-cold PBS, resuspended in ice-cold NT-2 Buffer (with 400 U/mL RNase inhibitor) and kept for 10 min on ice. 500 µL of lysates were sonicated to obtain RNA fragments (Bioruptor, 15 sec ON/45 sec OFF for 10 min at 4 °C on LOW settings). For both STAU1/2 CLIP, lysates were cleared by centrifugation for 5 min at 12,000 x g at 4 °C. Lysates (400 µg) were incubated for 16 h at 4 °C with primary antibodies in NT-2 Buffer (with 400 U/mL RNase inhibitor). Dynabeads™ Protein G (ThermoFisher) were added to the samples and incubated with gentle rotation for 2 h at 4 °C. Immunocomplexes were washed four times with NP-40 Buffer (20 mM Tris [pH 7.5], 100 mM KCl, 5 mM MgCl_2_, 0.5 % NP-40) and processed for RNA extraction/purification with GeneJet RNA purification kit (ThermoFisher) with the addition of Proteinase K treatment, according to the manufacturer’s protocol. Same volume of purified RNA samples was analyzed by RT-qPCR with sequence-specific primers (Table S1).

### Electrophoretic mobility shift assay (EMSA)

Nuclear extracts were incubated with double-stranded oligonucleotide probe containing a consensus C/EBP binding site sequence labeled with [γ-^32^P]ATP as previously described.^15^

### Quantitative real-time PCR

Total RNA was extracted and purified using QIAshredder (Qiagen) and GeneJet RNA purification kit (ThermoFisher) and reverse transcribed by using Maxima first strand cDNA synthesis kit for RT-qPCR with dsDNase (ThermoFisher) according to the manufacturers’ protocols. Relative gene expression was measured by quantitative PCR using SsoAdvanced SYBR Green Supermix (Bio-Rad) and mouse or human gene-specific primers (listed in Table S1). *Ppia/PPIA* or *B2m/B2M* were used as normalization reference genes.

### RNA FISH and immunofluorescence (IF) staining

For RNA FISH, Quanti-Gene ViewRNA ISH Cell Assay (ThermoFisher) was used according to the manufacturer’s protocol with the following modifications. Cells were plated at 1-2 × 10^4^ cells per chamber on µ-slides VI0.4 (Ibidi). After 1-2 days, cells were rinsed in ice-cold Cytoskeleton Buffer (CB; 10 mM MES pH 6.1, 150 mM NaCl, 5 mM MgCl_2_, 5 mM EGTA, and 5 mM glucose) ^45^, permeabilized and fixed in ice-cold pre-fixative mix in CB (4% paraformaldehyde, 0.01% glutaraldehyde, 0.05% saponin) ^46^ for 10 min at 4 °C. Cells were then incubated with ice-cold fixative mix in CB (4% paraformaldehyde, 0.01% glutaraldehyde) for 2 h at 4 °C. To better preserve localization of mRNA and proteins, cells were treated with SHIELD mix (SHIELD epoxy/SHIELD OFF buffer, 3:1 v/v) (Life Canvas Technology; Table S1) for 30 min at 37 °C ^47^. Cells were rinsed with CB twice and quenched with 50 mM NH_4_Cl (in CB) for 5 min at 20 °C. After FISH hybridization steps, cells were rinsed in PBS-S (0.05% Saponin, in PBS) and blocked with 5% albumin in PBS-S for 1 h at 20 °C. Cells were then incubated with the indicated primary antibodies (Table S1) for 16-18 h at 4 °C in 5% albumin in PBS-S. Cells were rinsed three times for 5 min at 20 °C with PBS-S and incubated for 1 h at 20 °C with AlexaFluor-conjugated secondary antibodies (Table S1) in 5% albumin in PBS-S. Cells were rinsed twice with PBS-S, co-stained with 0.1 mg/mL DAPI (in PBS) for 2 min at 20 °C, and then rinsed three times with PBS. Slides were kept in PBS at 4 °C for 1-3 days until imaging. Images were acquired using either a Zeiss LSM 780 or 880 confocal microscopes or a Zeiss LSM 880 AiryScan super-resolution microscope. Images were processed and analyzed using Zen black and Zen blue software (Zeiss).

### Live cell imaging

Cells were transduced with supernatants of lentiviral constructs expressing fluorescently tagged proteins. After selection with the appropriate marker, cells were plated at 2 × 10^4^ cells per chamber on µ-slides VI0.4 (Ibidi). The cells were observed by confocal microscopy the following day.

### Image analysis

#### Relative nuclear proximity index (RNPI)

RNPI is an ImageJ-based tool that measures the average distance of cytoplasmic signals (in this case *CEBPB* mRNA labeled by RNA FISH) to the nuclear boundary, per cell, normalized to a uniform distribution of the same signals. Details of the method have been described.^28^

#### mRNA distribution in cell areas

To quantify mRNA distribution in different areas of the cell, Imaris software version 9.8 (Bitplane, Belfast, UK) was used to fit mRNA signals to the spots module and protein signals to the surface module. The identified surfaces and spots were combined in the cells module to further elucidate the specific numbers of mRNA spots within a defined region of the cell. One spots module, three surface modules, and two cell modules were used to conduct the analysis. The mRNA signals were defined setting a defined size for the spot of 0.400 µm in XY and using a manual quality threshold value according to visual inspection of the image. There were multiple surfaces defined nuclei, kinase region, and cell boundaries. Nuclei were defined using the surface module and DAPI staining to identify individual nuclei. The protein signal from immunofluorescent staining of the CK2 kinase was used to define the kinase region using a manual threshold of high intensity signals and smoothing to find a defined region for CK2. The cell boundaries were determined by both oversaturating the kinase signal and using a DIC image of the cells. Two cell modules were built. Cell module one identified the cell boundary, nuclei, and mRNA spots. Cell module one was used to find mRNA spots in the cytoplasm and total mRNA spots within the entire cell. Cell module two identified the kinase region, nuclei, and mRNA spots. Cell module 2 was used to define mRNA spots in the kinase region. Number of *CEBPB* mRNA molecules per cell and % of *CEBPB* mRNA overlapping the kinase region per cell were calculated.

#### Nearest neighbor analysis

DiAna is an ImageJ plugin to measure distances between objects of two different species (such as object with two different fluorescence labels in optical microscopes images).^48^ In this application, DiAna is used to measure in the cytoplasm the shortest distance from one punctate species to another punctate species. The species were *CEBPB* transcripts labeled by RNA-FISH and CK2α foci labeled by immunofluorescence. Prior to executing DiAna, punctate signals in each channel were detected and the borders of the cytoplasm and nuclei for each cell were drawn. Punctate signals were detected by first choosing a minimum threshold intensity for puncta in the image. Then the puncta were detected by the 3D Maxima Finder plugin that is part of the 3D Suite in ImageJ. The settings for the Finder were the minimum threshold, an XY radius of 5, a Z radius of 1 and the noise level set to the square root of the minimum threshold. Only detected puncta inside the cytoplasm border and outside the nucleus border were used. Each cell in an image was analyzed separately. DiAna was implemented using the “Center-to-Center” and “Distance” options to measure the Euclidean distance from the centers of puncta of one species to the nearest center of puncta of the other species. Generally, the first punctate species was fewer in number than the other punctate species. Stepwise instructions and an ImageJ macro for executing the method will be provided upon request.

#### mRNA decay assay

mRNA decay rates were determine using 5-ethynyluridine (EU) labeled mRNA chase assay. A549 cells were plated at 10^5^ cells/well in 6-well dishes. After 24 h, 0.1 mM EU was added to the media for 16 h. After incubation, cells were either harvested (time 0) or media was replaced without EU and the cells were cultured for the indicated time points before RNA harvesting to allow degradation of the labeled RNA (chase). The isolated EU-labeled RNA was used with the Click-iT Nascent RNA Capture Kit (ThermoFisher) according to the manufacturer’s instructions.

### Proximity ligation assay (PLA)

A549 cells were plated at 4 × 10^4^ cells per chamber on µ-slides VI0.4 (Ibidi). The next day, cells were rinsed in ice-cold PBS and fixed with methanol for 15 min at −20 °C, permeabilized with 0.05% Saponin in PBS for 5 min at 20 °C and rinsed three times with PBS. After fixation, Duolink® in situ detection system (MilliporeSigma) was used according to the manufacturer’s instructions (volumes were adjusted accordingly for µ-slides). Images were acquired using a Zeiss LSM 780 confocal microscope.

### Colony forming assays

Cells were plated at 2000 cells per 100 mm culture dish. After 10-14 days in culture, colonies were fixed in 4% formaldehyde in PBS for 10 min, stained with 0.1% crystal violet for 30 min and counted.

### Focus formation assays

Cells were plated at 1000 cells per 100 mm culture dish with 1.5 × 10^5^ untransformed NIH 3T3 or *p19^Arf-/-^* MEFs (lawn). After 10-14 days, cells were fixed in 4% formaldehyde in PBS for 10 min, stained with 0.1% crystal violet for 30 min and foci were counted.

### Senescence-associated β-Galactosidase (SA-βGal) assays for cultured cells

Cells were plated at 2.5 × 10^4^ cells per well in 6-well plates. After 2-3 days, cells were stained using a senescence detection kit (MilliporeSigma) with modifications of the manufacturer’s instructions. For mouse cell lines, the pH of the staining solution mix was lowered by adding 1.54 µL of 2 M HCl per 1 mL.

### SA-βGal staining of tumor-bearing lung sections

Mice were euthanized following the approved ACUC protocol. Lung lobes were dissected and individually frozen in OCT on dry ice. Frozen sections were cut at 10 microns and stored at −80 °C until staining. Immediately prior to staining, slides were fixed in 1% formaldehyde/0.2% glutaraldehyde/0.2% Igepal for 10 min. Slides were then rinsed and washed in PBS for 10 minutes. Several drops of β-gal staining solution (1 mg/ml Xgal, 40 mM citric acid/Na phosphate buffer pH 5.8, 5 mM K_3_[Fe(CN)_6_], 5 mM K_4_Fe(CN)_6_, 150 mM NaCl, 2 mM MgCl_2_) were placed on the slides. The slides were then placed in a dark water bath at 37 °C overnight. After staining, slides were rinsed and washed in PBS for 10 minutes, followed by two 5 min washes in distilled water. Slides were counterstained with a 0.1% Neutral Red solution for 0.5 to 2 min. Finally, slides were dehydrated in 100% ethanol, cleared through four changes of xylene, mounted with Permount, and cover-slipped. All slides were digitally scanned at 20x magnification utilizing an Aperio AT2 whole-slide scanner. Lung nodules were annotated, and the percentage of positive cells was quantified using the HALO software, employing a cytonuclear algorithm and area quantification. Areas of artifact and background staining, such as those found in papillary nodules, were excluded from the analysis.

### Immunohistochemical (IHC) staining of tissue sections

For Ki67 IHC staining, slides generated from paraffin blocks (5 µm sections) were dewaxed using xylene and then hydrated using a series of graded ethyl alcohols. Slides were stained with Hematoxylin and Eosin-Y (H&E) following standard protocols using the Sakura® Tissue-Tek® Prisma™ automated stainer. A regressive staining method was used. This method intentionally overstains tissues and then uses a differentiation step (Clarifier/Bluing reagents) to remove excess stain. After staining was completed, the slides were coverslipped using the Sakura® Tissue-Tek™Glass® automatic coverslipper, dried and were ready for review. IHC staining was performed on LeicaBiosystems’ BondRX autostainer with the following conditions: Epitope Retrieval 1 (Citrate) for 20 min, Ki67 (Cell Signaling Technology #12202, 1:200 for 30 min), and the Bond Polymer Refine Detection Kit (LeicaBiosystems #DS9800, with omission of the PostPrimary Reagent). Rabbit monoclonal IgG XP® Isotype Control (Cell Signaling Technology) was used in place of Ki67 for the negative control. Slides were removed from the Bond autostainer, dehydrated through ethyl alcohols, cleared with xylenes, and coverslipped. H&E and IHC slides were scanned at 20X using an Aperio AT2 scanner (Leica Biosystems, Buffalo Grove, IL) into whole slide digital images.

For S100a9, CD206 and CD86 staining, FFPE blocks were sectioned at 5 microns in preparation for multiplex IHC. Staining was performed using a Leica Bond RX autostainer (Leica Biosystems). Initial antigen retrieval was performed using EDTA buffer for 20 min at 100 °C on the Bond autostainer. The primary antibody for S100a9 (Cell Signaling #73425) was diluted 1:500, with a 30-minute incubation time. S100a9 antibody detection was accomplished using the Bond Polymer Refine Detection kit (Leica Biosystems #DS9800) with the Post-Primary reagent, DAB, and Hematoxylin removed from Leica’s default staining protocol. Fluorescent Opal570 reporter dye (Akoya Biosciences # FP1488001KT) was added to the tissue per the manufacturer’s instructions. Following application of the Opal570 reagent, EDTA antigen retrieval solution was applied again to the tissues for 20 min at 95 °C. CD206 antibody (Cell Signaling #24595) was applied at a 1:400 dilution for 30 min. CD206 antibody detection was accomplished using the Bond Polymer Refine Detection kit (Leica Biosystems #DS9800) with the Post-Primary reagent, DAB, and Hematoxylin removed from Leica’s default staining protocol. Fluorescent Opal520 reporter dye (Akoya Biosciences # FP1487001KT) was added to the tissue per the manufacturer’s instructions. After application of the Opal520 reagent, EDTA antigen retrieval solution was applied again to the tissues for 20 min at 95 °C. CD86 antibody (Cell Signaling #19589) was applied at a 1:50 dilution for 60 min. CD86 antibody detection was accomplished using the Bond Polymer Refine Detection kit (Leica Biosystems #DS9800) with the Post-Primary reagent, DAB, and Hematoxylin removed from Leica’s default staining protocol. Fluorescent Opal690 reporter dye (Akoya Biosciences # FP1497001KT) was added to the tissue per the manufacturer’s instructions. All slides were digitally scanned at 20x using an Aperio FL whole-slide scanner. Normal mouse spleen was used as a positive control for all stains. The scanned slides were analyzed using HALO (Indica Labs, Inc.) Initially, benign and carcinomatous lesions were annotated separately. Subsequently, the percentage of positive cells was quantified using HighPlex FL (v3.2.2) algorithm.

### Image analysis of tissue samples

All image analysis was performed using HALO imaging analysis software (v3.3.2541.300; Indica Labs, Corrales, NM), and image annotations were performed in a blinded fashion. The lung sections were annotated with HALO manual annotation tools. Nonspecific regions such as esophagus, thymus, and trachea were excluded from analysis. Areas of artifact such as folds and tears were also omitted from analysis. The HALO random forest tissue classifier was used to automatically differentiate neoplastic cells from normal lung parenchyma to determine total tumor burdens. Ki67 image analysis was performed using cytonuclear algorithm in HALO version 3.3 to determine percentage of positive cells. HALO MiniNet AI tissue classifier was used to automatically differentiate tumor cells from normal lung parenchyma and to differentiate benign adenomas form malignant adenocarcinomas. The size of nuclei was measured from Ki67 IHC stained slides. Multiple nodules (adenomas and adenocarcinomas) were selected in two different layers. HALO Nuclei Seg (plugin) classifier was used to segment nuclei.

### Histological examination

In the *Kras^LA2^* model, expanding and replacing pulmonary parenchyma are multifocal, variably-sized, up to 7 mm diameter, well-demarcated, unencapsulated neoplastic foci composed of cuboidal to columnar epithelial cells arranged in either packets, solid sheets, and glands or forming papillary structures, supported by a fine fibrovascular stroma. Neoplastic cells in adenomatous nodules have variably-distinct cell borders, moderate amounts of eosinophilic cytoplasm, and round to oval nuclei with finely-stippled chromatin, and indistinct nucleoli. The mitotic rate is less than 1 per 20 high-power field (HPF). Multifocally, within one or more nodules, neoplastic cells forming adenocarcinomas display anisocytosis, which arranges in disorganized glands or solid sheets. Neoplastic cells have either scant to moderate amount of eosinophilic to amphophilic cytoplasm or have clear cytoplasm. Nuclei are moderately pleomorphic with fine to coarse chromatin and have prominent nucleoli. Mitotic rate is 0 to 2 per 20 HPF. Within the adjacent parenchyma, multifocally alveolar spaces often contain alveolar macrophages.

### Statistical analyses

Quantitative data are presented as means ± standard deviation with the number of measurements (“n”) indicated in the figure legends, representing biological replicates in cell culture experiments and the number of mice in animal studies. Statistical analysis was performed in GraphPad Prism using Student’s t test function, unless otherwise indicated in the figure legends. P-values less than 0.05 were considered significant.

## Notes

### Competing Interest Statement

The authors have declared no competing interest.

### Summary of Updates

This version of the manuscript has been revised to update certain figures with additional experimental replicates and certain experiments were removed. Further characterization of lung tumors in delGRE mice has been added to Fig. 7.

