## Supplementary Figures and legends and critical reagents list for "3’UTR-directed, kinase proximal mRNA decay inhibits C/EBPβ phosphorylation/activation to suppress senescence in tumor cells"

Figure S1

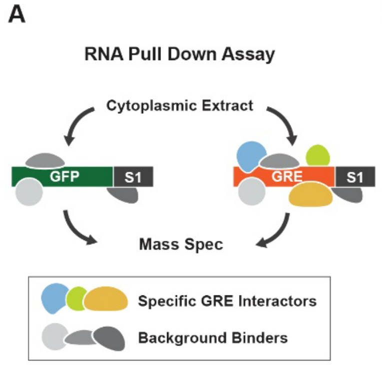

B

SASP mRNA Expression Relative to shCtrl Vector

| Knockdown | *Il6* | | *Il1a* | | *Cxcl1* | | *Ccr1* | |
|---|---|---|---|---|---|---|---|---|
| | $\beta^{UTR}$ | shβ | $\beta^{UTR}$ | shβ | $\beta^{UTR}$ | shβ | $\beta^{UTR}$ | shβ |
| Upf1 | 300.6 | 15.4 | 490.3 | 11.0 | 62.9 | 13.2 | 21.5 | 3.8 |
| Strbp | 22.0 | 0.6 | 108.8 | 1.0 | 65.4 | 3.3 | 6.7 | 1.8 |
| Elavl1 | 6.8 | 0.6 | 0.7 | 0.8 | 3.2 | 5.2 | 29.8 | 1.4 |
| Ilf3 | 3.7 | 2.8 | 4.3 | 1.2 | 0.7 | 0.5 | 1.5 | 0.6 |
| Igf2bp3 | 3.0 | 1.2 | 4.8 | 0.9 | 3.4 | 2.8 | 0.9 | 0.2 |
| Fubp3 | 2.7 | 0.5 | 3.6 | 0.7 | 4.3 | 4.2 | 11.8 | 2.4 |
| Anxa1 | 2.3 | 0.9 | 1.3 | 0.5 | 2.6 | 1.5 | 8.9 | 0.6 |
| Igf2bp1 | 1.8 | 0.8 | 8.1 | 0.6 | 1.6 | 0.6 | 0.9 | 0.3 |
| Mapre1 | 1.6 | 0.3 | 2.5 | 0.0 | 0.9 | 0.4 | 0.8 | 0.2 |
| Igf2bp2 | 1.5 | 0.3 | 17.4 | 0.3 | 1.6 | 0.7 | 0.9 | 0.2 |
| Hnrnpd | 1.4 | 0.7 | 1.5 | 0.1 | 1.0 | 0.7 | 0.6 | 0.3 |
| Dhx9 | 1.3 | 0.3 | 1.4 | 0.1 | 1.4 | 0.8 | 1.4 | 0.4 |
| Celf1 | 1.2 | 0.5 | 6.2 | 1.7 | 1.7 | 1.5 | 7.9 | 0.9 |
| Fmr1 | 1.0 | 0.4 | 0.7 | 0.0 | 0.9 | 0.6 | 0.7 | 0.3 |
| Hnrnpa2b1 | 0.8 | 0.5 | 0.4 | 0.1 | 1.2 | 0.7 | 1.1 | 0.4 |
| Anxa2 | 0.8 | 1.0 | 1.1 | 1.0 | 1.6 | 1.0 | 2.3 | 1.0 |
| Plec | 0.8 | 0.3 | 2.3 | 0.5 | 1.4 | 2.2 | 3.6 | 0.7 |
| Nono | 0.5 | 0.2 | 1.1 | 0.3 | 1.2 | 1.2 | 0.5 | 0.1 |
| Fubp2 | 0.5 | 0.2 | 1.3 | 0.6 | 1.1 | 3.9 | 2.3 | 0.5 |
| Fubp1 | 0.4 | 0.3 | 2.2 | 1.0 | 1.6 | 2.4 | 4.3 | 1.3 |
| Ilf2 | 0.3 | 0.2 | 0.5 | 0.0 | 0.5 | 0.1 | 4.2 | 1.7 |

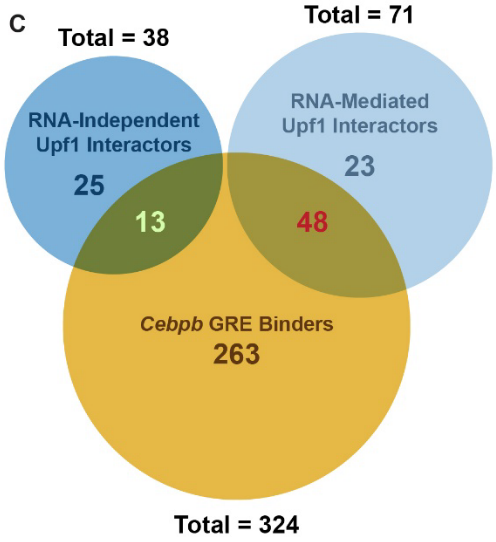

**Figure S1. Upf1 interacts with the *Cebpb* GRE to inhibit C/EBPβ-dependent SASP gene expression, Related to Figure 1.**

**(A)** An RNA affinity purification scheme to identify novel GRE-interacting proteins. The procedure is based on the method of Butter *et al.*[1] **(B)** Screening of selected C/EBPβ UPA candidate genes. Each gene was ablated in NIH 3T3$^{RAS}$ cells overexpressing *Cebpb*$^{UTR}$ (βUTR) or depleted for C/EBPβ (shβ), and expression of proinflammatory SASP mRNAs was measured by

RT-qPCR. **(C)** Overlap between GRE binding proteins and known RNA-dependent and RNA-independent UPF1 interactors.[2]

Figure S2

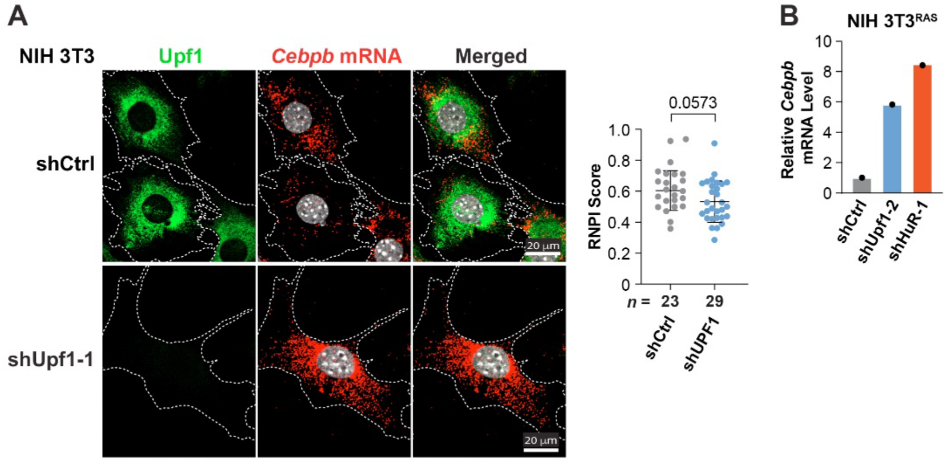

**Figure S2. Silencing of UPF1 increases the perinuclear abundance of *CEBPB* mRNAs, Related to Figure 2.**

**(A)** Upf1 knockdown in NIH 3T3 cells increases the perinuclear segregation of *Cebpb* transcripts. *Cebpb* mRNA was detected using single molecule RNA FISH together with immunostaining for Upf1. Right: a relative nuclear proximity index (RNPI) score[3] was determined for each cell. *n* = number of cells analyzed. **(B)** Upf1 depletion increases total *Cebpb* mRNA levels in NIH 3T3$^{RAS}$ cells. mRNA levels were determined by RT-qPCR analysis, normalized to *Ppia* mRNA.

Figure S3

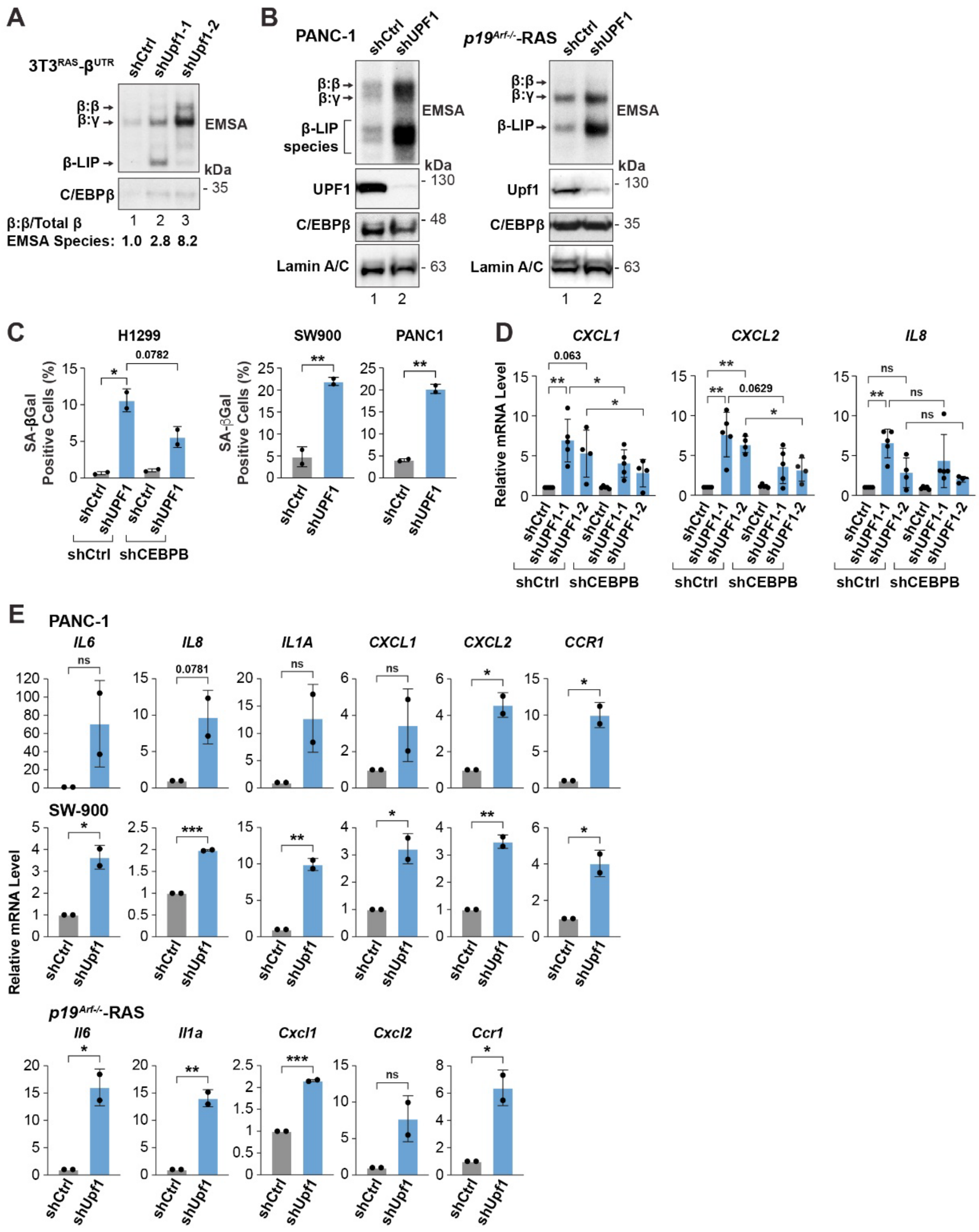
A
3T3RAS-βUTR
shCtrl
shUpf1-1
shUpf1-2
β:β
β:γ
β-LIP
EMSA
kDa
C/EBPβ
35
β:β/Total β
EMSA Species:
1 2 3
1.0 2.8 8.2
B
PANC-1
shCtrl
shUPF1
β:β
β:γ
β-LIP species
EMSA
kDa
UPF1
130
C/EBPβ
48
Lamin A/C
63
1 2
p19Arf-/--RAS
shCtrl
shUPF1
β:β
β:γ
β-LIP
EMSA
kDa
Upf1
130
C/EBPβ
35
Lamin A/C
63
1 2
C
H1299
SW900
PANC1
SA-βGal Positive Cells (%)
shCtrl
shUPF1
shCtrl
shCEBPB
0.0782
D
CXCL1
CXCL2
IL8
Relative mRNA Level
0.063
0.0629
ns
shCtrl
shUPF1-1
shUPF1-2
shCtrl
shCEBPB
E
PANC-1
IL6
IL8
IL1A
CXCL1
CXCL2
CCR1
0.0781
ns
SW-900
Relative mRNA Level
shCtrl
shUpf1
p19Arf-/--RAS
Il6
Il1a
Cxcl1
Cxcl2
Ccr1

**Figure S3. UPF1 depletion activates the pro-senescence functions of C/EBPβ in tumor cells, Related to Figure 3.**

**(A)** Upf1 silencing stimulates C/EBPβ DNA-binding activity in $3T3^{RAS}$-$\beta^{UTR}$ cells. Nuclear extracts were analyzed by EMSA using a C/EBP binding site probe. The C/EBPβ homodimer EMSA band (β:β) was quantitated and normalized to total C/EBPβ binding (all species). **(B)** Depletion of UPF1 stimulates C/EBPβ DNA-binding activity in PANC-1 cells and $HRAS^{G12V}$-transformed $p19^{Arf-/-}$ MEFs. **(C)** Analysis of senescence (SA-βGal staining) in a panel of human tumor cell lines depleted for UPF1. **(D)** Activation of SASP genes in UPF1-depleted A549 cells, without and with *CEBPB* knockdown. The indicated genes were analyzed by RT-qPCR. **(E)** Effect of UPF1 silencing on SASP gene expression in a subset of the cell lines shown in **(C)** and $HRAS^{G12V}$-transformed $p19^{Arf-/-}$ MEFs. The indicated genes were analyzed by RT-qPCR.

Statistical significance was determined using Student's t test. *$p < 0.05$,**$p < 0.01$,***$p < 0.001$; ns, not significant.

### Figure S4

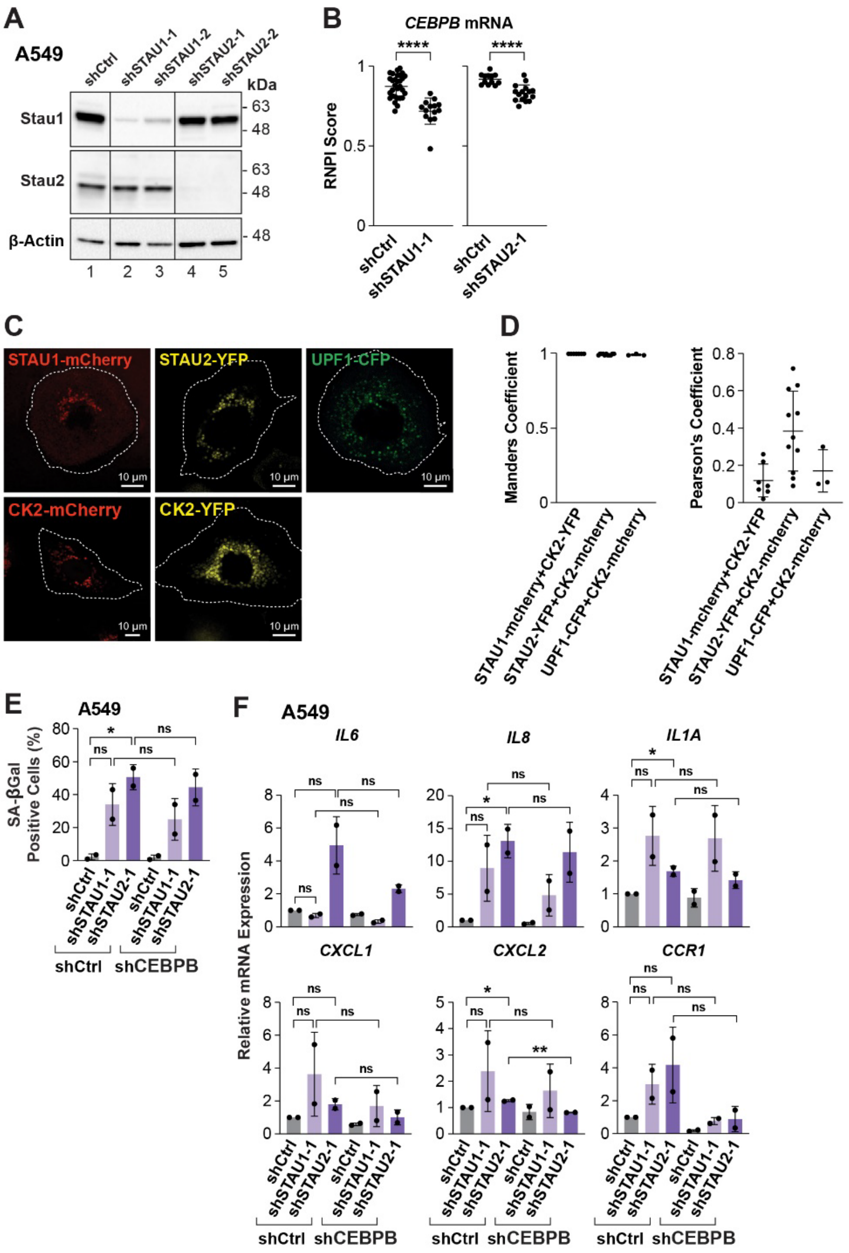
A
A549
shCtrl
shSTAU1-1
shSTAU1-2
shSTAU2-1
shSTAU2-2
kDa
Stau1
Stau2
β-Actin
63
48
1 2 3 4 5
B
CEBPB mRNA
RNPI Score
****
shCtrl
shSTAU1-1
shSTAU2-1
C
STAU1-mCherry
STAU2-YFP
UPF1-CFP
CK2-mCherry
CK2-YFP
10 µm
D
Manders Coefficient
Pearson's Coefficient
STAU1-mcherry+CK2-YFP
STAU2-YFP+CK2-mcherry
UPF1-CFP+CK2-mcherry
E
A549
SA-βGal Positive Cells (%)
ns
*
shCtrl
shCEBPB
F
A549
IL6
IL8
IL1A
CXCL1
CXCL2
CCR1
Relative mRNA Expression
**

**Figure S4. STAU depletion in tumor cells induces senescence and weakly activates SASP genes, Related to Figure 4.**

**(A)** STAU1 and STAU2 knockdown efficiencies in A549 cells as assessed by immunoblotting. **(B)** Relative nuclear proximity index (RNPI) scores for *CEBPB* mRNA FISH signals in A549 cells, without and with STAU1/2 depletion (analysis of experiment in Fig. 4D). **(C)** Images of fluorescently tagged STAU1, STAU2, UPF1 and CK2α expressed individually in A549 cells. STAU1, STAU2 and UPF1 display perinuclear localization, demonstrating their intrinsic perinuclear targeting independent of co-expressed CK2α. **(D)** Manders overlap and Pearson's correlation coefficients for fluorescently tagged STAU1, STAU2 and UPF1 co-expressed with CK2α (analysis of experiment in Fig. 4F). **(E)** Analysis of senescence (SA-βGal staining) in control or *CEBPB*-depleted A549 cells without and with STAU1/2 depletion ($n \geq 400$ cells). **(F)** STAU1 or STAU2 depletion in A549 cells modestly upregulates SASP genes in a manner partially dependent on C/EBPβ.

Statistical significance was determined using Student's t test. *$p < 0.05$, ****$p < 0.0001$; ns, not significant.

### Figure S5

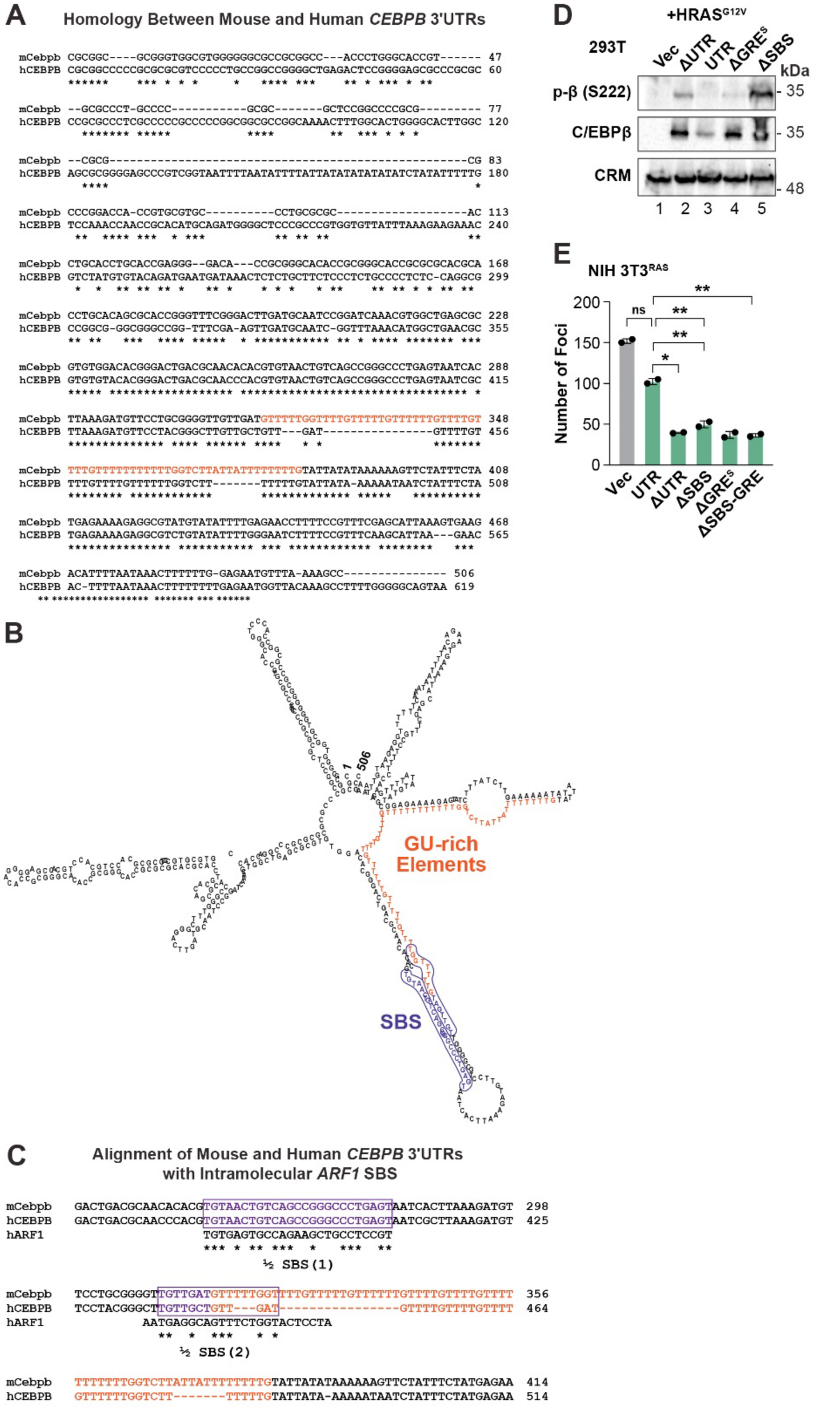
A
Homology Between Mouse and Human CEBPB 3'UTRs
B
GU-rich Elements
SBS
C
Alignment of Mouse and Human CEBPB 3'UTRs with Intramolecular ARF1 SBS
½ SBS (1)
½ SBS (2)
D
+HRAS G12V
293T
Vec
ΔUTR
UTR
ΔGRE S
ΔSBS
kDa
p-β (S222)
C/EBPβ
CRM
- 35
- 35
- 48
1 2 3 4 5
E
NIH 3T3 RAS
Number of Foci
ns
Vec
UTR
ΔUTR
ΔSBS
ΔGRE S
ΔSBS-GRE

**Figure S5. Conservation of the SBS/GRE region in mouse and human *CEBPB* 3'UTRs and analysis of SBS/GRE mutants, Related to Figure 5.**

**(A**) Alignment of mouse and human *CEBPB* 3'UTRs. The strongest conservation occurs in the 3' regions surrounding the GU-rich sequence (orange). **(B)** An RNA secondary structure prediction model (RNAFold) shows stem-loops formed by part of the GU-rich sequence (orange) paired with a 5' adjacent sequence containing a putative SBS (purple). **(C)** A predicted STAU binding site (SBS) adjacent to the GRE region is identified by sequence homology with an intramolecular SBS identified in the *ARF1* 3'UTR[4]. Diagram shows conservation of the potential SBS element (purple) in mouse and human *CEBPB* and its position relative to the GRE (orange). Note the partial overlap between the GRE and SBS elements. **(D)** $HRAS^{G12V}$-induced phosphorylation on C/EBPβ Ser222 (CK2 site) in 293T cells, comparing several 3'UTR deletion mutants. **(E)** Focus formation assays for NIH $3T3^{RAS}$ cells expressing *Cebpb* 3'UTR mutant constructs.

Statistical significance was determined using Student's t test. *$p < 0.05$, **$p < 0.01$; ns, not significant.

Figure S6

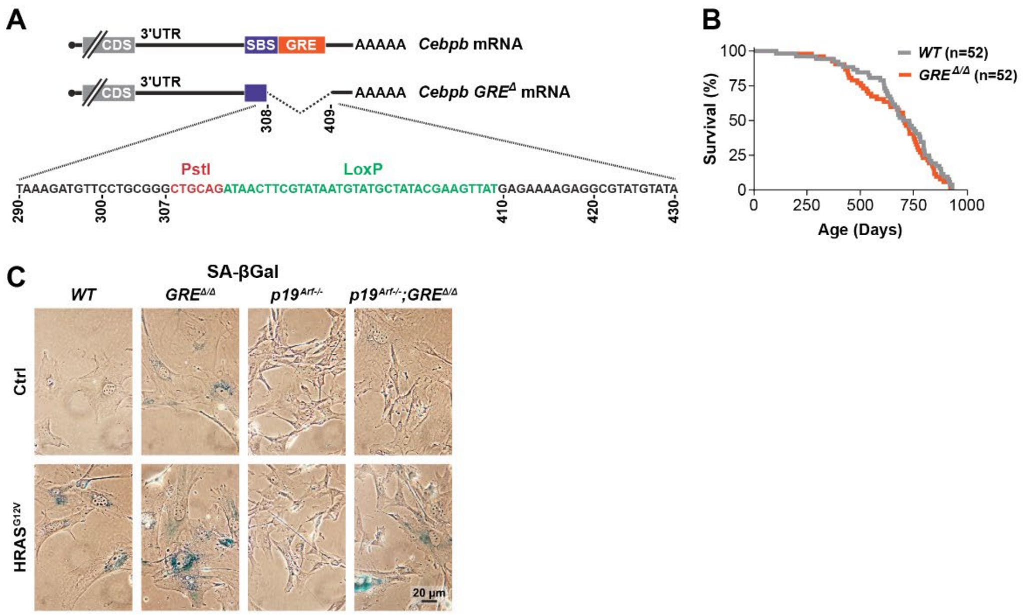

**Figure S6. RAS induces senescence in *Cebpb-GRE*$^{\Delta/\Delta}$ MEFs lacking the tumor suppressor p19$^{Arf}$, Related to Figure 6.**

**(A)** Diagram of the mouse *Cebpb* gene and *GRE*$^{\Delta}$ allele. The *GRE*$^{\Delta}$ allele was generated by Cre-mediated recombination of a floxxed allele containing LoxP sites flanking the GRE motif. **(B)** Kaplan-Meier survival curves of *WT* and *GRE*$^{\Delta/\Delta}$ mice, showing no significant difference in life spans (Log-rank test; $p = 0.35$). *n* = number of animals. **(C)** Representative images of SA-βGal staining for *WT*, *GRE*$^{\Delta/\Delta}$, *p19*$^{Arf-/-}$, and *p19*$^{Arf-/-}$*;GRE*$^{\Delta/\Delta}$ MEFs, without or with HRAS$^{G12V}$ expression. Quantification of SA-βGal positive cells in several independently-derived MEF lines is shown in Fig. 6C.

Figure S7

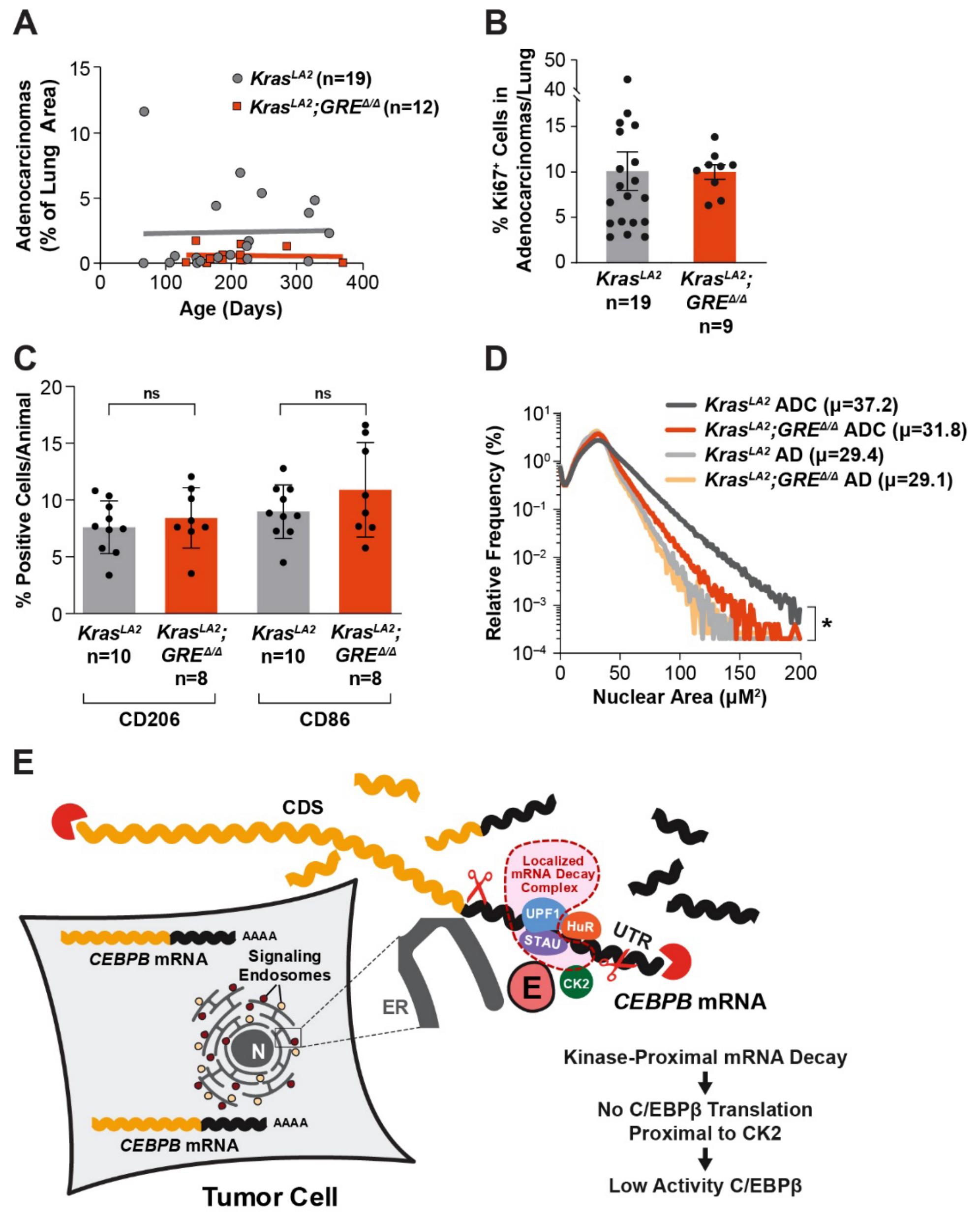

**Figure S7. Analysis of tumors in *Kras$^{LA2/+}$;Cebpb-GRE$^{\Delta/\Delta}$* mice and summary model for kinase proximal mRNA decay, Related to Figure 7.**

**(A)** Lack of correlation between age at clinical sacrifice and adenocarcinoma burdens in *Kras*$^{LA2/+}$and *Kras*$^{LA2/+}$*;GRE*$^{\Delta/\Delta}$ mice. The adenocarcinoma data of Fig. 7C is presented as a scatter plot versus age. **(B)** Percent Ki67 positive cells within adenocarcinoma lesions for each genotype. **(C)** Macrophage infiltration in *Kras*$^{LA2/+}$and *Kras*$^{LA2/+}$*;GRE*$^{\Delta/\Delta}$ tumors. Lung sections were stained for CD206 (M2-type) or CD86 (M1-type) macrophages; data are shown as % positive cells in tumor areas per animal. **(D)** Frequency distribution ($log_{10}$ scale) of nuclear size (area) in adenomas (AD) and adenocarcinomas (ADC) of both genotypes. Population means (μ) are shown. The total number of nuclei analyzed by HALO are: *Kras*$^{LA2}$ ADC (n = 1,834,153); *Kras*$^{LA2}$*;GRE*$^{\Delta/\Delta}$ ADC (n = 498,806); *Kras*$^{LA2}$ AD (n = 494,181); *Kras* $^{LA2}$*;GRE*$^{\Delta/\Delta}$ AD (n = 380,307). **(E)** Model depicting 3'UTR-directed, kinase proximal *CEBPB* mRNA decay in tumor cells. *CEBPB* mRNA decay occurs at perinuclear CK2α-containing signaling endosomes through STAU proteins localized to these organelles. Signaling endosomes in tumor cells are embedded with the perinuclear ER network.[3] Localized mRNA decay also requires UPF1, which could be recruited to endosomes by STAU proteins.[4] Degradation of *CEBPB* transcripts at $CK2^+$ endosomes prevents the protein from becoming phosphorylated by CK2, maintaining C/EBPβ in a low activity state that facilitates senescence bypass.

#### Table S1.

| REAGENT/RESOURCE | SOURCE | CATALOG NUMBER/SEQUENCE/REFERENCE |
|---|---|---|
| **Antibodies** | | |
| β-Actin | Santa Cruz Biotechnology | sc-1616, sc-47778 |
| Anti-mouse AlexaFluor-488 | ThermoFisher Scientific | A21202 |
| Anti-mouse AlexaFluor-568 | ThermoFisher Scientific | A10037 |
| Anti-mouse AlexaFluor-647 | ThermoFisher Scientific | A21236 |
| Anti-mouse horseradish peroxidase-conjugated antibody | Promega | W4028 |
| Anti-rabbit AlexaFluor-488 | Abcam | ab150077 |
| Anti-rabbit AlexaFluor-647 | Abcam | ab150075 |
| Anti-rabbit horseradish peroxidase-conjugated antibody | Promega | W4018 |
| C/EBPβ | Abcam | ab32358 |
| C/EBPβ | NSJ Bioreagents | R31588 |
| phospho-C/EBPβ (S223); rat | In house | |
| CSNK2A1 (CK2α) | Abcam | ab10466 |
| CSNK2A1 (CK2α) | Santa Cruz Biotechnology | sc-365762 |
| GFP | Santa Cruz Biotechnology | sc-8334 |
| HuR | Santa Cruz Biotechnology | sc-5261 |
| Lamin A/C | Cell Signaling Technology | 2032 |
| SMG1 | Santa Cruz Biotechnology | sc-374557 |
| SMG5 | ProteinTech | 12694-I-AP |
| SMG6 | ThermoFisher Scientific | PA5-60165 |
| SMG7 | GeneTex | GTX121320 |
| STAU1 | Abcam | ab137100 |

| | | |
|---|---|---|
| STAU2 | Sigma | HPA019155 |
| α-Tubulin | Santa Cruz Biotechnology | sc-8035 |
| UPF1 | Cell Signaling Technology | 12040 |
| UPF1 | Santa Cruz Biotechnology | sc-393594 |
| phospho-UPF1 (Ser1127) | MilliporeSigma | 07-1016 |
| **Bacterial and virus strains** | | |
| DH5α | ThermoFisher Scientific | 18265017 |
| Stbl3 | ThermoFisher Scientific | C737303 |
| **Chemicals, peptides, and recombinant proteins** | | |
| 25 cm Acclaim PepMap C18 column | ThermoFisher Scientific | Easy nLC 1200 |
| Actinomycin D | MilliporeSigma | A9415 |
| Dynabeads® MyOne™ Streptavidin C1 | ThermoFisher Scientific | 65001 |
| Dynabeads™ Protein G for Immunoprecipitation | ThermoFisher Scientific | 10004D |
| Micrococcal Nuclease (MNase) | ThermoFisher Scientific | 88216 |
| μ-Slide VI 0.4 | Ibidi | 80606 |
| 4X Laemmli sample buffer | Bio-Rad Laboratories | 1610747 |
| Phosphatase Inhibitor Cocktail Set I | MilliporeSigma | 524624 |
| Phosphatase Inhibitor Cocktail Set II | MilliporeSigma | 524625 |
| Protease Inhibitor Cocktail Set I | MilliporeSigma | 539131 |
| Protein A, HRP conjugate | MilliporeSigma | 18-160 |
| SYBR Green | Bio-Rad Laboratories | 1725275 |
| SUPERase-In RNase Inhibitor (20 U/μL) | ThermoFisher Scientific | AM2696 |
| RNase A | ThermoFisher Scientific | R1253 |

| | | |
|---|---|---|
| yeast tRNA | ThermoFisher Scientific | AM7119 |
| **Critical commercial assays** | | |
| Click-iT™ Nascent RNA Capture Kit | ThermoFisher Scientific | C10365 |
| Duolink® In Situ PLA® Probe Anti-Rabbit PLUS | MilliporeSigma | DUO92002-30RXN |
| Duolink® In Situ PLA® Probe Anti-Mouse MINUS | MilliporeSigma | DUO92004-30RXN |
| Duolink® In Situ Detection Reagents Green | MilliporeSigma | DUO92014-30RXN |
| Duolink® In Situ Wash Buffers, Fluorescence | MilliporeSigma | DUO82049-4L |
| GeneJet RNA purification kit | ThermoFisher Scientific | K0732 |
| Maxima First Strand cDNA Synthesis Kit for RT-qPCR, with dsDNase | ThermoFisher Scientific | K1672 |
| QiaShredder | Qiagen | 79656 |
| Quanti-Gene ViewRNA ISH Cell Assay | ThermoFisher Scientific | QVC0001 |
| SHIELD | LifeCanvas Technology | 250 mL |
| SuperSignal™ West Dura Extended Duration Substrate | ThermoFisher Scientific | 34076 |
| SuperSignal™ West Pico PLUS Extended Duration Substrate | ThermoFisher Scientific | 34579 |
| TranscriptAid T7 High Yield Transcription Kit | ThermoFisher Scientific | K0441 |
| **Experimental models: cell lines** | | |
| NIH 3T3 | ATCC | CRL-1658 |
| A549 | ATCC | CCL-185 |
| PANC-1 | FNL/RAS Initiative | |
| SW-900 | FNL/RAS Initiative | |
| IMR-90 | ATCC | CCL-186 |

| | | |
|---|---|---|
| HEK 293T | ATCC | CRL-3216 |
| **Experimental models: organisms/strains** | | |
| Mouse: C57Bl/6Ncr | The Jackson Lab | MGI:2160593 |
| Mouse: 129/SV | The Jackson Lab | MGI:2161069 |
| Mouse: p19$^{Arf-/-}$ (CdKn2a-Arf) B6.129 | NCI-Frederick Repository | |
| Mouse: ACTB-Cre C57Bl/6 | Mark Lewandoski Lab (NCI) | [5] |
| Mouse: Kras$^{G12D-LA2/+}$ 129/SV | Esta Sterneck Lab (NCI) | [6] |
| **Oligonucleotides** | | |
| T7-GRE PCR primers | IDT | 5'-CGTTAATACGACTCACTATAGGGTTGTTGATGTTTTTGGTTTTGTTT |
| Apt-GRE PCR primers | IDT | 5'-CATGGCCCGGCCCGCGACTATCTTACGCACTTGCATGATTCTGGTCGGTCCCATGGATCCATAGAAATAGAACTTTTTTA |
| T7-GFP PCR primers | IDT | 5'-CGTTAATACGACTCACTATAGGCTTTCCAAAATGTCGTAACAACTCC; [1] |
| Apt-GFP PCR primers | IDT | 5'-CATGGCCCGGCCCGCGACTATCTTACGCACTTGCATGATTCTGGTCGGTCCCATGGATCCCTTCAGGGTCAGCTTGCCG; [1] |
| Human B2M qPCR primers | Qiagen | QT00088935 |
| Human CCL2 qPCR primers | Qiagen | QT00212730 |
| Human CCR1 qPCR primers | Qiagen | QT00047740 |
| Human CEBPB qPCR primers | Qiagen | QT00998494 |
| Human CXCL1 qPCR primers | Qiagen | QT00199752 |
| Human CXCL2 qPCR primers | Qiagen | QT00013104 |

| | | |
|---|---|---|
| Human CXCL5 qPCR primers | Qiagen | QT00203686 |
| Human IL1A qPCR primers | Qiagen | QT00001127 |
| Human IL6 qPCR primers | Qiagen | QT00083720 |
| Human IL8 qPCR primers | Qiagen | QT00000322 |
| Human PPIA qPCR primers | Qiagen | QT01866137 |
| Human S100A9 qPCR primers | Qiagen | QT00018739 |
| Mouse B2m qPCR primers | Qiagen | QT01149547 |
| Mouse Ccl2 qPCR primers | Qiagen | QT00167832 |
| Mouse Ccr1 qPCR primers | Bio-Rad Laboratories | qMmuCID0006862 |
| Mouse Cebpb (CDS) qPCR primers | Qiagen | QT00320313 |
| Mouse Cebpb1 (UTR) qPCR forward primers | IDT | CGCACGCACCTGCACA |
| Mouse Cebpb1 (UTR) qPCR reverse primers | IDT | GTTACACGTGTGTTGCGTCAG |
| Mouse Cebpb2 (UTR) qPCR forward primers | IDT | CAATCCGGATCAAACGTGGC |
| Mouse Cebpb2 (UTR) qPCR reverse primers | IDT | CCGCAGGAACATCTTTAAGTGA |
| Mouse Cebpb3 (UTR) qPCR forward primers | IDT | CTGACGCAACACACGTGTAAC |
| Mouse Cebpb3 (UTR) qPCR reverse primers | IDT | AAACAAAACCAAAAACATCAACAACC |
| Mouse Cebpb4 (UTR) qPCR forward primers | IDT | TCACTTAAAGATGTTCCTGCGG |
| Mouse Cebpb4 (UTR) qPCR reverse primers | IDT | CGCCTCTTTTCTCATAGAAATAGAAC |
| Mouse Cebpb5 (UTR) qPCR forward primers | IDT | GTTCTATTTCTATGAGAAAAGAGGCG |
| Mouse Cebpb5 (UTR) qPCR reverse primers | IDT | GGCTTTTAAACATTCTCCAAAAAAGTTTATTAA |
| Mouse Cxcl1 qPCR primers | Qiagen | QT00115647 |
| Mouse Cxcl2 qPCR primers | Bio-Rad Laboratories | qMmuCED0003897 |
| Mouse Cxcl5 qPCR primers | Bio-Rad Laboratories | qMmuCED0047657 |
| Mouse Dhx9 qpCR primers | Bio-Rad Laboratories | qMmuCID0023309 |
| Mouse Il1a qPCR primers | Qiagen | QT00113505 |

| | | |
|---|---|---|
| Mouse Il6 qPCR primers | Qiagen | QT00098875 |
| Mouse Ppia qPCR primers | Qiagen | QT00247709 |
| Mouse S100a9 qPCR primers | Qiagen | QT00105252 |
| Human CEBPB ViewRNA Cell Probe Set AlexaFluor-546 | ThermoFisher Scientific | VX-06/VA1-18129 |
| Mouse Cebpb ViewRNA Cell Probe Set AlexaFluor-546 | ThermoFisher Scientific | VX-06/VB1-10094-VC |
| EMSA probe (C/EBP binding site sequence is underlined) | IDT | 5'-GATCCATATCCCTGATTGCGCAATAGGCTCAAAA; [7] |
| Mouse *Cebpb* GRE$^{flox}$ allele | IDT | LoxP sites flanking *Cebpb GRE* region (UTR nucleotides 308-409, underlined). PstI site inserted upstream of 5'LoxP and BclI site inserted upstream of 3'LoxP: CTGCAGATAACTTCGTATAATGTATGCTATACGAAGTTATGTTGTTGATGTTTTTGGTTTTGTTTTTGTTTTTTGTTTTGTTTTGTTTTTTTTTTTGGTCTTATTATTTTTTTTGTATTATATAAAAAAGTTCTATTTCTATGATCATAACTTCGTATAATGTATGCTATACGAAGTTAT |
| **Recombinant DNA** | | |
| pBabe (puro) | Addgene | 1764 |
| pBabe$\beta^{UTR}$ (puro) | In house | [8] |
| pBabe$\beta^{\Delta UTR}$ (puro) | In house | [9] |
| pBabe$\beta^{UTR}$-*Malat1*-Scrbl (puro) | In house | 5'-CCTGGTGATGTGTTGTGTTCTTGTACTCTGATAGCTTCGTGCTGCTGTTGTATCTTACTATAGTGTTGGCATCGGCACCCATGTCATGGTTAGTATTCTGATCGCTCAGGTCCTTAGATAGGATGCCTTGCACGAGACGTGATACAATCTTCGTTCTAATACAATCAGAAGATA |
| pBabe$\beta^{UTR}$-*Malat1* (puro) | In house | 5'-GATTCGTCAGTAGGGTTGTAAAGGTTTTTCTTTTCCTGAGAAAACAACCTTTTGTTTTCTCAGGTTTTGCTTTTTGGCCTTTCCCTAGCTTTAAAAAAAAAAAAGCAAAAGACGCTGGTGGCTGGCACTCCTGGTTTCCAGGACGGGGTTCAAGTCCCTGCGGTGTCTTTGCTT |
| pBabe$\beta^{\Delta SBS}$ (puro) | In house | Deletion on βUTR nucleotides 264-316, replaced by BamHI |
| pBabe$\beta^{\Delta GRE\text{-}S}$ (puro) | In house | Deletion on βUTR nucleotides 317-382, replaced by BamHI |

| | | |
|---|---|---|
| pBabeβ$^{\Delta SBS\text{-}GRE}$ (puro) | In house | Deletion on βUTR nucleotides 264-382, replaced by BamHI |
| pSirenRetroQ-puro | Clontech/Takara | |
| pSirenRetroQ-puro mouse shAnxa1 | In house | AGCAACCATCATTGACATTCT |
| pSirenRetroQ-puro mouse shAnxa2 | In house | CTTCGATGCTGAGAGGGATGC |
| pSirenRetroQ-puro mouse shCelf1 | In house | GAGCCAACCTGTTCATCTA |
| pSirenRetroQ-puro mouse shDhx9 | In house | GCCAGAGACTTTGTTAACTATT; [10] |
| pSirenRetroQ-puro mouse shFmr1 | In house | GCATGTGATGCTACGTATA |
| pSirenRetroQ-puro mouse shFubp1 | In house | ATACAGATAGCACCTGATAGT |
| pSirenRetroQ-puro mouse shFubp2 | In house | GGGACACCATGATCTGAATGT |
| pSirenRetroQ-puro mouse shFubp3 | In house | GGATTAGTCCAGAAAGAGC |
| pSirenRetroQ-puro mouse shHnrnpa2b1 | In house | CGTGCTGTAGCAAGAGAGG; [11] |
| pSirenRetroQ-puro mouse shHnrnpd | In house | GACGCCAGTAAGAACGAGG |
| pSirenRetroQ-puro mouse shIgf2bp1 | In house | CCGGGAGCAGACCAGGCAA; [12] |
| pSirenRetroQ-puro mouse shIgf2bp2 | In house | GGCATCAGTTTGAGGACTA; [13] |
| pSirenRetroQ-puro mouse shIgf2bp3 | In house | GGGAAGAATTTATGGAAAA |
| pSirenRetroQ-puro mouse shIlf2 | In house | CCATTTGGACATCAAGGTG |
| pSirenRetroQ-puro mouse shIlf3 | In house | GGACGGACAGAAGTTTCAAGG |
| pSirenRetroQ-puro mouse shMapre1 | In house | GAAGAAAGTGAAATTCCAAGC |
| pSirenRetroQ-puro mouse shNono | In house | GACCTTTACACAGCGTAGC; [14] |
| pSirenRetroQ-puro mouse shPlec | In house | GGCAGGACCGTGACCATCT |
| pSirenRetroQ-puro mouse shStrbp | In house | GTCTATAGGTACTTGTAAT |
| pSirenRetroQ-puro mouse shUpf1 | In house | GATGCAGTTCCGTTCCATC |

| | | |
|---|---|---|
| pSIRIP-puro shRNA p19Arf-2 | Addgene | 14090 |
| pSuperRetro-neo | OligoEngine | |
| pSuperRetro-neo shCebpb-1 | In house | GATGTTCCTGCGGGGTTGT; [9] |
| pSuperRetro-neo shCEBPB | In house | GAAGAAACGTCTATGTGTA; [9] |
| pSuperRetro-neo shELAVL1 | In house | GAGGCAATTACCAGTTTCA; [8] |
| CMV13p>control (hygro) | Protein Expression Lab (FNL) | 10449-M03-665 |
| CMV51p>Hs.HRAS G12V | Protein Expression Lab (FNL) | R700-M18-665 |
| pMD2.G | Addgene | 12259 |
| pMDLg/pRRE | Addgene | 12251 |
| pRSV/Rev | Addgene | 12253 |
| pReceiver-Lv123-CSNK2A1-eYFP (puro) | Genecopoeia | EX-T0147-Lv123 |
| pReceiver-Lv216-Csnk2a1-mCherry (puro) | Genecopoeia | EX-Mm01978-Lv216 |
| pReceiver-Lv155-STAU1-mCherry (neo) | Genecopoeia | EX-U1331-Lv155 |
| pReceiver-Lv123-STAU2-eYFP (puro) | Genecopoeia | EX-Z7308-Lv123 |
| pReceiver-Lv130-STAU2-mCherry (puro) | Genecopoeia | EX-Z7308-Lv130 |
| pReceiver-Lv127-UPF1-eCFP (puro) | Genecopoeia | EX-Z0991-Lv127 |
| pReceiver-Lv130-UPF1-mCherry (puro) | Genecopoeia | EX-Z0991-Lv130 |
| MISSION® pLKO.1-puro eGFP shRNA Control Plasmid | MilliporeSigma | SHC005 |
| MISSION® pLKO.1-puro ELAVL1 shRNA Plasmid (1) | MilliporeSigma | TRCN0000276186 |
| MISSION® pLKO.1-puro ELAVL1 shRNA Plasmid (2) | MilliporeSigma | TRCN0000276129 |
| MISSION® pLKO.1-puro SMG1 shRNA Plasmid (1) | MilliporeSigma | TRCN0000196274 |
| MISSION® pLKO.1-puro SMG1 shRNA Plasmid (2) | MilliporeSigma | TRCN0000244939 |
| MISSION® pLKO.1-puro SMG1 shRNA Plasmid (3) | MilliporeSigma | TRCN0000197107 |

| | | |
|---|---|---|
| MISSION® pLKO.1-puro SMG1 shRNA Plasmid (4) | MilliporeSigma | TRCN0000244937 |
| MISSION® pLKO.1-puro SMG5 shRNA Plasmid (1) | MilliporeSigma | TRCN0000421114 |
| MISSION® pLKO.1-puro SMG5 shRNA Plasmid (2) | MilliporeSigma | TRCN0000130800 |
| MISSION® pLKO.1-puro SMG6 shRNA Plasmid (1) | MilliporeSigma | TRCN0000040014 |
| MISSION® pLKO.1-puro SMG6 shRNA Plasmid (2) | MilliporeSigma | TRCN0000419520 |
| MISSION® pLKO.1-puro SMG7 shRNA Plasmid (1) | MilliporeSigma | TRCN0000292264 |
| MISSION® pLKO.1-puro SMG7 shRNA Plasmid (2) | MilliporeSigma | TRCN0000292266 |
| MISSION® pLKO.1-puro STAU1 shRNA Plasmid (1) | MilliporeSigma | TRCN0000159875 |
| MISSION® pLKO.1-puro STAU1 shRNA Plasmid (2) | MilliporeSigma | TRCN0000164920 |
| MISSION® pLKO.1-puro STAU2 shRNA Plasmid (1) | MilliporeSigma | TRCN0000219963 |
| MISSION® pLKO.1-puro STAU2 shRNA Plasmid (2) | MilliporeSigma | TRCN0000102356 |
| MISSION® pLKO.1-puro UPF1 shRNA Plasmid (1) | MilliporeSigma | TRCN0000413098 |
| MISSION® pLKO.1-puro UPF1 shRNA Plasmid (2) | MilliporeSigma | TRCN0000022254 |
| MISSION® pLKO.1-puro UPF1 shRNA Plasmid (3) | MilliporeSigma | TRCN0000022257 |
| MISSION® pLKO.1-puro UPF1 shRNA Plasmid (4) | MilliporeSigma | TRCN0000022255 |
| MISSION® pLKO.1-puro Upf1 shRNA Plasmid (1) | MilliporeSigma | TRCN0000274484 |
| MISSION® pLKO.1-puro Upf1 shRNA Plasmid (2) | MilliporeSigma | TRCN0000009664 |
| **Software and algorithms** | | |
| Biorad CFX Manager | Bio-Rad Laboratories | 184500 |
| DAVID (2021 Release) | LHRI, FNL | [15] |
| GraphPad Prism 9.2.0 | GraphPad Software | https://www.graphpad.com/scientific-software/prism/ |
| HALO imaging analysis v3.3.2541.300 | Indica Labs | |

| | | |
|---|---|---|
| HALO cytonuclear algorithm | Indica Labs | |
| HALO random forest tissue classifier | Indica Labs | |
| HALO MiniNet AI tissue classifier | Indica Labs | |
| HALO nuclei seg (plugin) classifier | Indica Labs | |
| Imaris | Oxford Instruments | |
| ImageJ | NIH | [16] |
| Leica | Leica Biosystems | |
| SEQUEST and the Percolator validator algorithms in Proteome Discoverer 2.2 | ThermoFisher Scientific | |
| Zen black | Zeiss | |
| Zen blue | Zeiss | |
